# TimeTeller: a New Tool for Precision Circadian Medicine and Cancer Prognosis

**DOI:** 10.1101/622050

**Authors:** Denise Vlachou, Georg A. Bjarnason, Sylvie Giacchetti, Francis Lévi, David A. Rand

## Abstract

Recent studies have established that the circadian clock influences onset, progression and therapeutic outcomes in a number of diseases including cancer and heart disease. Therefore, there is a need for tools to measure the functional state of the circadian clock and its downstream targets in patients. We provide such a tool and demonstrate its clinical relevance by an application to breast cancer where we find a strong link between survival and our measure of clock dysfunction. We use a machine-learning approach and construct an algorithm called TimeTeller which uses the multi-dimensional state of the genes in a transcriptomics analysis of a single biological sample to assess the level of circadian clock dysfunction. We demonstrate how this can distinguish healthy from malignant tissues and demonstrate that the molecular clock dysfunction metric is a potentially new prognostic and predictive breast cancer biomarker that is independent of the main established prognostic factors.

## Introduction

The cell-endogenous circadian clock regulates tissue-specific gene expression in cells which drives rhythmic daily variation in metabolic, endocrine, and behavioural functions. Indeed, around half of all mammalian genes are expressed with a 24-hour rhythm (*3, 4*). Moreover, recent studies demonstrated that the circadian clock influences therapeutic outcomes in a number of diseases including heart disease and cancer (*5–11*), and that disruption of the normal circadian rhythm and sleep (e.g. through shift work) is associated with higher risk of obesity, hypertension, diabetes, chronic heart disease, stroke and cancer (*12–15*).

A principal aim of circadian medicine (*17, 18*) is to develop techniques and methods to integrate the relevance of biological time into clinical practice. However, it is difficult to monitor the functional state of the circadian clock and its downstream targets in patients. Consequently, there is a critical need for tools to do this that are practical in a clinical context. Our focus is on the development of such a technique. We present a machine-learning approach to measuring circadian clock functionality from the expression levels of 10-16 key genes in a single tissue sample.

To illustrate its utility, our algorithm, TimeTeller, is applied to breast cancer where previous studies have highlighted the relevance of circadian clocks for carcinogenesis and treatment effects (*19–22*) but where no simple method would currently allow its measurement in daily oncology practice. We find a strong association between overall and disease-free survival and our measure of clock dysfunction in breast cancer patients.

Our strategy in designing a dysfunction metric is to try and estimate the conditional distributions *P*(*t*|*g*) of the time *t* conditional on the simultaneous expression *g* = (*g*_1_,…, *g*_*G*_) of a set of *G* clock or clock-associated genes where *G* ≈10 – 16. Then the time of collection of an independent sample can be estimated as the time *T* of the maximum of *P*(*t*|*g*_*S*_) where *g*_*S*_ is the expression vector for this sample. Ideas from statistics and information theory enable us to estimate the confidence interval for this maximum likelihood estimator (MLE) *T* for any given degree of confidence using the likelihood ratio function Λ_*g*_ (*t*) = *P*(*g*|*t*) / *P*(*g*|*T*) which TimeTeller estimates. If this confidence interval is small then our MLE *T* is certain and we regard this as indicating a clock providing precise timing and having good functionality. A large interval is similarly associated with imprecise timing and dysfunctionality. Our metric Θ is associated with the length of such a confidence interval (Note S4) and therefore increasing Θ is associated with worse dysfunctionality. Consequently, we call Θ a *clock dysfunction metric*.

There are now several published algorithms which aim to estimate the time at which a transcriptomic dataset was collected using the expression levels of the core clock genes (*17, 23–28*). While these have been used to study aspects of clock dysfunction by calculating a measure for a dataset as a whole and comparing a diseased dataset with a healthy one (*24, 26*), they are not designed to measure clock dysfunction in single samples. The ideas from statistics and information theory mentioned above show that the key quantities determining Θ are the covariance structure of *P*(*g*|*t*) and the way that *g*_*S*_ moves as time *t* advances (Note S4). Therefore, to measure precision, an algorithm must address these quantities and none of the existing algorithms has been designed to do this (Notes S10.4 & S13.2, Fig. S5E,F). The availability of a single sample algorithm such as TimeTeller potentially allows the determination of clock dysfunction in an individual patient along the course of his or her disease and treatments, the study of its relations with patient outcomes, and the use of such measurements in clinical practice.

The circadian timing system controls several critical molecular pathways for cancer processes and treatment effects. While in the cells of most healthy tissues the cell cycle is gated or phase-locked by the circadian clock (*29, 30*), cancer cells often escape this control and display altered molecular clocks (*31–33*). Dysregulation of clock genes promotes tumorigenesis (*25*) through mechanisms involving the cell cycle (*34, 35*), DNA damage (*36*), and metabolism (*37*). Moreover, the circadian clock rhythmically controls many molecular pathways which are responsible for large time-of-day dependent changes in drug toxicity and efficacy, whilst clock gene expression is correlated to anti-cancer drug sensitivity in cancer cell lines (*5, 6, 38, 39*). Altered expression of clock genes is associated with key oncogenic pathways and patient survival, and the correlation between clock genes and other genes is altered in cancerous tissues (*39*). Cadenas *et al.* (*40*) showed that the expression level of some circadian clock genes were associated with metastasis-free survival (MFS) in breast cancer, and discussed the relation with current prognostic factors. They also related gene expression levels (at a single time point that could differ between patients) to clock dysfunction at the dataset level. They did this by comparing the correlation between the expression of pairs of clock genes for the dataset associated with each prognostic stratum. Shilts *et al.* (*26*) developed two metrics based on comparing how the pattern of Spearman correlations between the clock genes differed in normal and cancerous tissue to conclude that circadian clock progression was perturbed in a range of human cancers. However, Cadenas *et al.* and Shilts *et al.* compared datasets as a whole and Shilts *et al.* further pointed out that their approach did not lend itself to assessing clock function in single samples.

In view of these results, it is of interest to apply TimeTeller’s ability to measure the functionality of the clock in an individual patient’s tumour tissue to determine whether this is a prognostic factor for treatment response and survival. We demonstrate here that TimeTeller can reliably measure dysfunction in single samples and that the dysfunction metric is an independent prognostic factor for both disease-free survival (DFS) and overall survival (OS) in a large cohort of patients with primary breast cancer.

It is important to note that our approach is very different from the common technique of clustering patients using gene expression from a candidate prognostic gene set asking for specific over- or under-expression of the genes in this set. Instead, from a single sample we directly assess the multi-dimensional state of the selected clock-associated genes and study the coordinated behaviour of all the genes together rather than focus on each gene separately. We are not looking for under- or over-expression per se but whether or not these rhythmic genes stand in the correct relationship with each other. Using this multidimensional technique we can measure the functionality of the clock system as a whole much more effectively (Note S4). Moreover, we avoid the observed tendency whereby many gene sets (including “random” ones) appear to be associated to breast cancer outcomes because of strong correlation with an organising signature such as meta-PCNA (*41*). Indeed, there is essentially no correlation between PCNA and our dysfunction metric (Note S10.5).

## Results

The mouse and human versions of TimeTeller are trained on two different datasets. The human training data set from Bjarnason *et al.* ((*42*), Note S5.1) comes from punch biopsies of oral mucosa taken every four hours over 24 hours from five females and five males. This human dataset was chosen because a key initial aim was to develop TimeTeller in order to analyse clock function in the tumour biopsies from the REMAGUS trial (*43*) and our analysis suggested that it was important to match the microarray technologies (Note S9.2) which in this case was Affymetrix U133 2.0. The mouse dataset, that we use is from Zhang *et al.* ((*3*), Note S5.1) and consists of the transcriptomes of 8 mouse tissues measured every 2 hours over 48 hours. The procedure for using these datasets to successfully produce TimeTeller’s probability model and likelihood functions is explained in Notes S1–3.

To determine the panel of genes for TimeTeller both the rhythmicity and synchronicity were analysed (Note S7, Table S1). This analysis is essential to ensure the choice of a panel of genes with good circadian rhythmicity combined with minimal variation across the relevant tissues and datasets. It typically produces a panel of between 10 and 16 gene probes and, for the human training dataset, the genes selected are all core clock genes or key clock-controlled genes, including *ARNTL* (BMAL1), *NPAS2*, *PER1*, *PER2*, *PER3*, *NR1D1* (REV-ERBα), *NR1D2* (REV-ERBβ), *CIART* (CHRONO), *TEF* and *DBP*. We call the vector consisting of the combined expression level of these genes the *rhythmic expression profile*.

TimeTeller works on the rhythmic expression profile *g* and estimates (up to a proportionality constant) an approximation *L*_*g*_(*t*) of the likelihood curve *P*(*t*|*g*) discussed above, which for Wild Type (WT) tissue should express the probability that the expression profile *g* was measured at time *t*. In our context this likelihood function is the primary mathematical object expressing the information about clock functionality. To correlate such measurements with clinical disease outcomes such as survival it is necessary to attempt to find associated scalar quantities that summarise the relevant aspects of the likelihood function. The metric Θ discussed above is one way to do this which is naturally suggested by statistics and information theory but there are likely to be other interesting metrics (such as the maximum value of *P*(*g*|*t*) as *t* varies). It depends upon three parameters. one of which, *l*_thresh_. is used to facilitate the consolidation of the local likelihood curves into a global one (Note S1.2, Figs. S1 & S6), while the other two, *ε* and *η*, are involved in the definition of Θ (Note S2). When comparing the values of Θ across different datasets it is important to use the same probability model and therefore the same values for these parameters.

We have tested the likelihood ratio functions (LRFs) and this clock dysfunction metric using both simulated and real data. Simulated data were obtained by developing a stochastic version of a relatively detailed published model of the mammalian circadian clock (*44*) and stochastically simulating this (Note S12). This data was used to design the algorithm and to test the effectiveness of TimeTeller, for example, to determine the advantage of local approaches over a global one (Figs. S1 & S6), and to analyse TimeTeller’s effectiveness in detecting the efficiency of partial knockdowns of various efficiencies of the central clock gene *BMAL1* (ARNTL). We found (Fig. S4A) that the efficiency of the knockdown was effectively recapitulated by an increase in Θ. We then applied TimeTeller to a number of mouse and human datasets.

### Training data provides a range of lower Θ values which is consistent across multiple datasets from healthy tissues

To evaluate TimeTeller’s likelihood curves *L*_*g*_(*t*), and the corresponding dysfunction metric Θ we firstly tested it on the training datasets using a leave-one-out approach. In particular, we asked if there is a threshold value Θ_GCF_ of Θ such that Θ < Θ_GCF_ can reliably be associated with the good clock function (GCF) of healthy or WT tissue. For the human Bjarnason *et al.* training data we removed the individuals one at a time, constructed the probability model for TimeTeller using the expression profiles from the other individuals and then used TimeTeller to estimate the times *T* and dysfunction Θ of the samples for the removed individual. For the mouse Zhang *et al.* data we carried out a similar leave-one-out approach but where a tissue rather than an individual was left out (Fig. S3).

The range of Θ values obtained are shown in the histogram in Fig. 3A,B and Fig. S3. For the human training set with *l*_thresh_ = −12 all samples have Θ < 0.1. For this reason and because of the shape of the tail of the histogram around 0.1, it is natural to define Θ_GCF_ = 0.1. Similar consideration lead to a choice of Θ_GCF_ = 0.225 for the mouse training set when *l*_thresh_ = −5 or −7. We cannot directly compare the mouse and human Θs because they use different training data and probability models.

We also see that the distributions of Θ values for a given individual (Bjarason *et al.* data) or tissue (Zhang *et al.* data) using the probability model determined by the other individuals or tissues are consistent across individuals and tissues (Fig. 2B & S9) and in the GCF range. We also analysed a number of other WT and healthy datasets and these uniformly have Θs in the GCF range (Figs. 3, S9, S12-14, S23-24). This supports the notion that the probability model can be used across multiple tissues.

**Figure 1.**
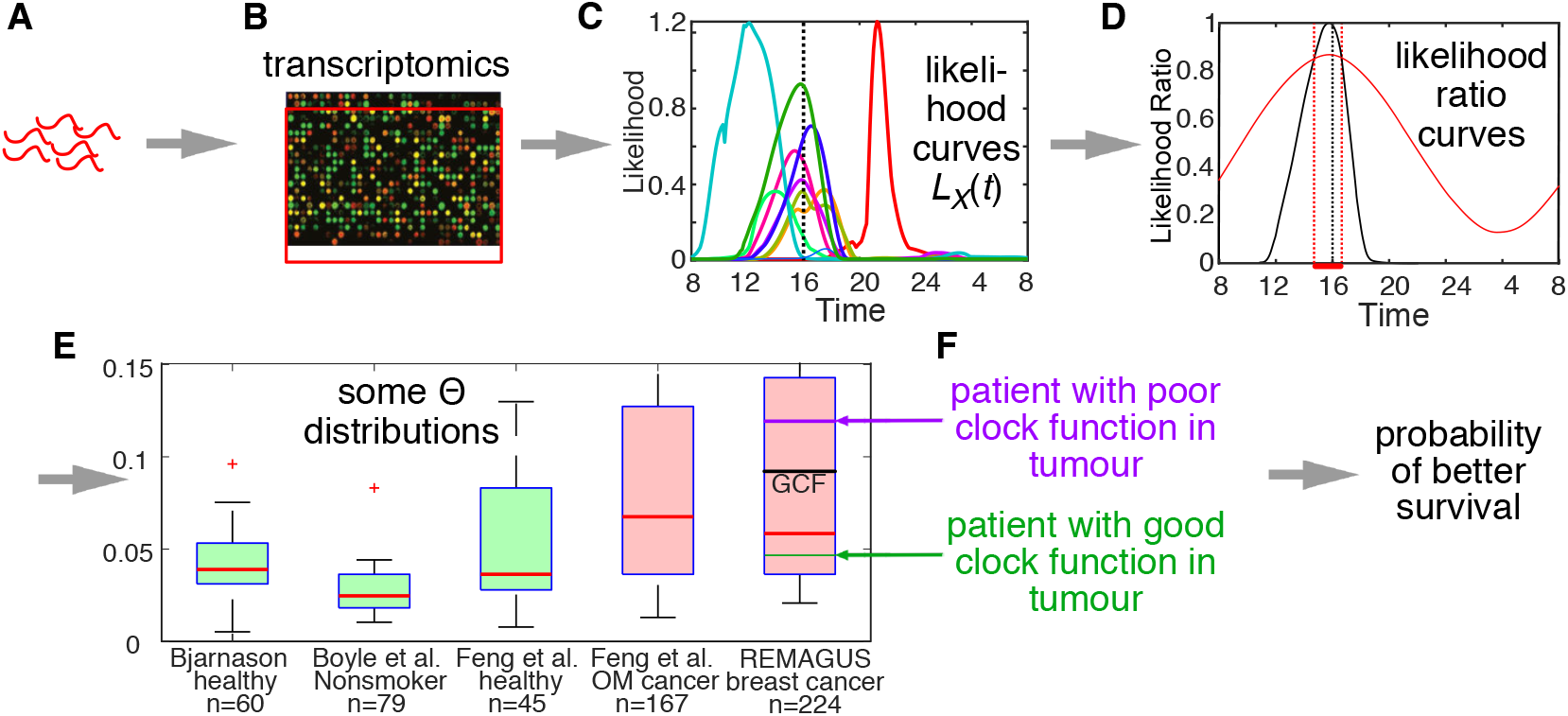
Graphical abstract. (A-E) From a tissue sample via the TimeTeller algorithm to an estimate of the rhythmic functionality of the clock for an individual. (F) Linking this estimate for breast cancer tumour clocks to survival.

**Figure 2.**
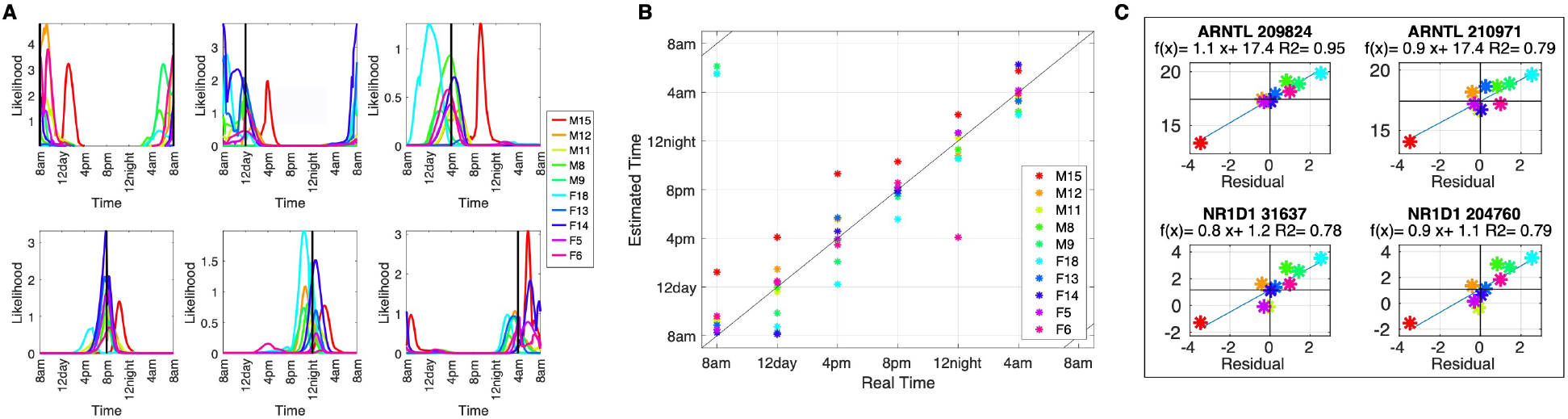
Likelihood curves and timing for human training data. (A) The likelihood functions *L*_*g*_(*t*) obtained for the samples from six of the times tested in the human training dataset. The actual time of each samples is indicated by a black vertical line. The maximum point of the likelihood functions is used to estimate the time when the sample was taken. In this and the other subfigures a leave-one-individual-out approach is used and the parameter *l*_thresh_ = −12. The 5 male (M) and 5 female (F) subjects are identified with a different colour consistently across all subfigures. (B) Scatter plots showing actual versus estimated time for the human training data ((41), Note S5.1). (B). For each individual the residual (estimated time - real time) is plotted against the phase of the gene of interest as measured by COSINOR and a linear fit to this is shown. See Fig. S3 for similar plots for all genes in the TimeTeller panel.

We verified that this range of GCF Θ values was associated with healthy timekeeping. The results are shown in Figs. 2, S9, S11 and S23-24. For the Zhang *et al.* mouse data and the Bjarnason *et al.* human data the mean absolute errors are 1.32 h and 1.36 h, respectively. However, in assessing accuracy we need to take account of the fact that there may be variation in clock gene phase across individuals and tissues. An analysis of the human training data shows that much of this apparent error is due to coherent phase variation in the clock genes possibly due to genetic and/or environmental factors. For example, Male15 and Female18 in Fig. 2B and 2D have consistent, yet opposite, phase shifts in their estimated times. To further understand this, we tested for a linear relation between the phase variation of the genes in our panel and apparent error. We plotted the error against the phase of each of the genes in the TimeTeller panel (Fig. 2D & Fig. S3), using COSINOR (*45*) to measure gene expression phase. For all the probes used we observed a significant linear relationship between apparent error and the variation in gene phase and are able to identify coherent phase variation in the clock genes for each individual across all genes in the rhythmic expression profile. Correcting for the phase of the genes gives a mean residual error of 0.56h (Fig. S3F). This corrected error is comparable with that for the best performing mouse tissue using TimeTeller or ZeitZeiger (*24, 26*). We hypothesise that low error is due to the fact that in the Bjarnason *et al.* data the samples for different times come from the same individual and tissue. These results suggest that if the real clock time of the sample is known by combining the observation of a Θ suggesting good clock function with an advanced or retarded time prediction. TimeTeller can identify coherent phase variation in an individual’s clock genes from a single sample.

For the mouse training data the use of the full 48-hour space allows us to observe that TimeTeller’s transcriptomic time signature at CT*t* is essentially the same as at CT(*t*+24) (Fig. S3). This means that there is no significant change to the circadian clock gene shape after the mice have been in the dark for an extra 24 hours.

Similar levels of timing accuracy with Θs in the GCF range were found in two mouse datasets from Le Martelot *et al*. (*46*) and Barclay *et al.* (*47*) (Fig. S4F-H). Timing for some diseased examples and its relation to second peaks is discussed below.

### TimeTeller can identify perturbed but functioning clocks

The gene *Nr1d1* (Rev-erbα) is regarded as the main controller of the Zeitgeber time (ZT)18-24 phase of the mammalian circadian clock (*1*) and knocking out *Nr1d1* leaves a functional but perturbed clock when compared to WT mice (*1*). Therefore, we applied TimeTeller to an experimental dataset Fang *et al.* (*1*) comparing liver samples of *Nr1d1* KO and WT mice entrained to light-dark (LD)12:12 cycles. Since *NR1D1* is one of the panel of genes used in TimeTeller it would not be surprising that TimeTeller could distinguish *Nr1d1* KO mice from WT mice, and indeed this is the case. Therefore, for this validation, we use a version of TimeTeller that excludes *Nr1d1* from the expression profile genes. This modified TimeTeller clearly detects that the *Nr1d1* KO mice have a functional but significantly perturbed clock when compared to WT mice (Fig. 3C). Although the WT (blue) likelihoods are wide, they produce relatively accurate and consistent estimations of ZT around 36h, with corresponding Θs in the good clock range and a mean absolute error of around 2 hours for time estimations of the WT data. This slightly raised estimation error, but good Θ values, could be explained by the discrepancy arising from the use of mice in constant darkness to train TimeTeller to estimate the time of mice that have been in regular LD cycles. The (red) KO likelihoods appear almost entirely flat if not plotted on a logarithmic scale (Fig. 3C). Another example where TimeTeller identifies perturbed human clocks is the study by Boyle *et al.* (*16*) of the effects of smoking on the oral mucosal transcriptome (Note 5.3) as the median Θ for smokers is significantly raised relative to non-smokers (Fig. S4B,C). Similarly, TimeTeller distinguishes the Θ distribution for samples from the liver of WT and sleep deprived mice ((*47*), Note S5.2 & Fig. S4G-H).

**Figure 3.**
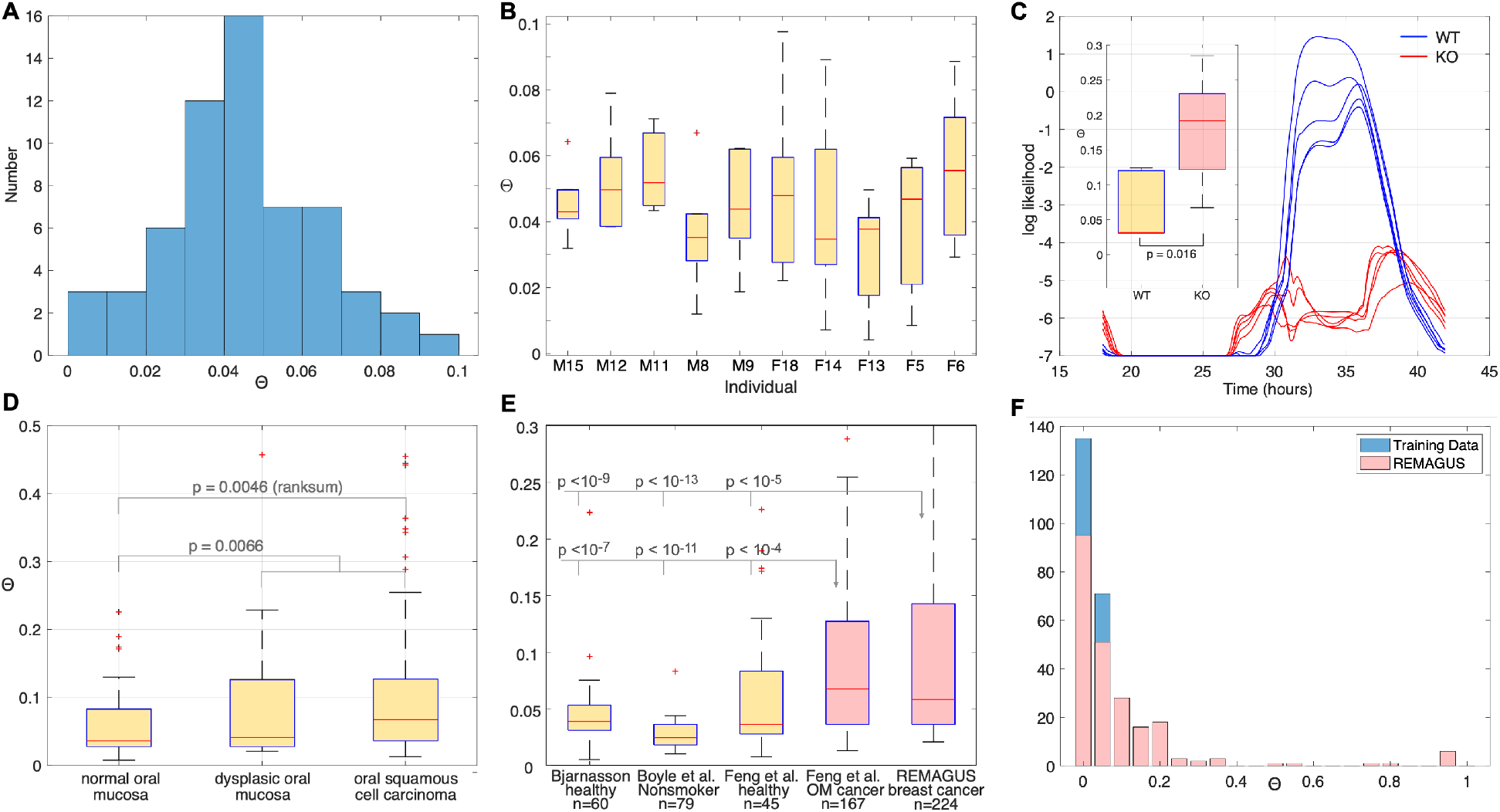
Distributions of Θ values for human datasets. For human datasets *l*_thresh_ = −12 and 15 probes are used. **A.** Histogram showing the distribution of Θ values for the human training data. **B.** Distributions of the Θ values (median, interquartiles and extremes) for each of the 10 individuals (M, males; F, females; with corresponding code number). **C**. Log likelihood functions for the MLE time *T* of mouse samples of the Fang *et al.* data using *l*_thresh_ = −7. They have been plotted on a logarithmic scale to reveal structure in the very low *Nr1d1* knockout (KO) curves. Blue curves represent WT samples with clear peaks around CT32-36, and the negligible amplitude curves in red for the KO data. C(inset). Box plot of Θ values for WT and *Nr1d1* KO data from (*1*). There is a significant difference between the groups. Wilcoxon’s logrank test p = 0.0159. **D.** The distribution of Θ values by disease group for the data on healthy or dysplastic oral mucosa and oral squamous cell carcinoma (OSCC) from Feng *et al*. (*2*). There is a highly significant difference in median values between the cancer group (n = 167) and both the normal mucosa group (n = 45) and the combined normal/dysplasia group n = 65), with the p-values indicated. **E.** Box plots of Θ distribution for healthy and cancerous tissue. As indicated they are from Bjarnason *et al.* (Note S5.1), Boyle *et al.* ((*16*), Note S5.3), Feng *et al.* ((*2*), Note S5.3), and the REMAGUS trial (Note S5.3). **F.** Histogram comparing the distribution of Θ values for healthy oral mucosa training data (blue) and breast cancer samples (red) from the REMAGUS clinical trial. The histograms are stacked.

### Healthy and malignant human tissue have different Θ distributions

In Fig. 3 we show the Θ distributions for a number of datasets. These include healthy datasets which served as controls in the indicated studies and our training datasets, and we observe consistent Θ distribution across all of these. The control Θ distributions can then be compared with two cancer datasets that used the same microarray technology, including oral squamous cell carcinoma from Feng *et al.* (41) and breast carcinomas from the REMAGUS trial discussed below (*43, 48–50*). Similarly, in Fig. 3(E,F) we compare the histogram of Θ values from the training data with that from the patients in the REMAGUS trial. For the human data we use *l*_thresh_ = −10 or −12. The results presented are for *l*_thresh_ = −12 but the results for *l*_thresh_ = −10 are entirely consistent. We observe that, although around half of the cancer data has Θ values in the same range as the healthy Bjarnason *et al.* data, the distributions for the cancer data are significantly biased towards larger values of Θ.

### The clock dysfunction metric Θ is associated with prognosis in breast cancer

To assess TimeTeller’s potential relevance, we used it to analyse tumour molecular clock dysfunction in the breast cancer patients who participated to the REMAGUS study. This multicenter randomised phase II clinical trial aimed to assess the response of primary breast cancer to different protocols of neoadjuvant chemotherapy according to tumour hormonal receptor status and HER2 expression (*43, 48–50*). Of the trial’s 340 patients, 226 had a pretreatment cancer biopsy using the same RNA extraction procedure and analysed with Affymetrix U133A microarrays. The patients’ characteristics are described in Table S2. With *l*_thresh_ = −12 thirteen of the samples have flat regions in the likelihood curves that contribute significantly to their value of Θ because the maximum value of their likelihood curve is very small (Fig. S2). Such Θs are not reliable and therefore our study population consists of the 209 patients with both ten-year survival data and reliable Θ computation (92.4% of the patients with tumour transcriptome determinations before the application of neoadjuvant chemotherapy). While the cause of death for four patients is unknown, for the others it was breast cancer progression.

To study whether Θ was indicative of survival we used the Cox proportional hazards model (*51*) to calculate the hazard ratios (HRs) associated with overall survival (OS) and disease free survival (DFS) for the whole study population and for the classical prognostic strata. There is a clear statistically significant survival advantage in both OS and DFS for patients with a Θ above the median value (Figs. 4A,D). However, since Θ is a continuous variable it is more natural instead to treat it as such and to ask whether a change in Θ is associated with a change in survival. In particular, this avoids having to choose a particular threshold for good clock function. In the multivariate continuous variable context, the meaning of the hazard rate HR is that if we hold the other prognostic factors constant at their mean HR values, the ratio of the hazard of the an individual with Θ’ to that of one with Θ is HR^(Θ^’ ^−Θ)^. Thus, HR is a measure of the effect on survival resulting from a change from Θ to Θ’ and if HR < 1 then the hazard decreases with increasing Θ. Since we want to compare the HR for Θ with the other prognostic factors which are categorized and given by 0 and 1 while Θ varies over a smaller range, for the comparison in Fig. 4 and Table S3, we scale Θ by 10 and use the modified hazard rate HR^1/10^ which is the hazard rate for 10×Θ or equivalently the ratio for a Θ increase of 0.1. Of course, the p-value for the alternative hypothesis HR < 1 is the same as for HR^1/10^ < 1.

**Figure 4.**
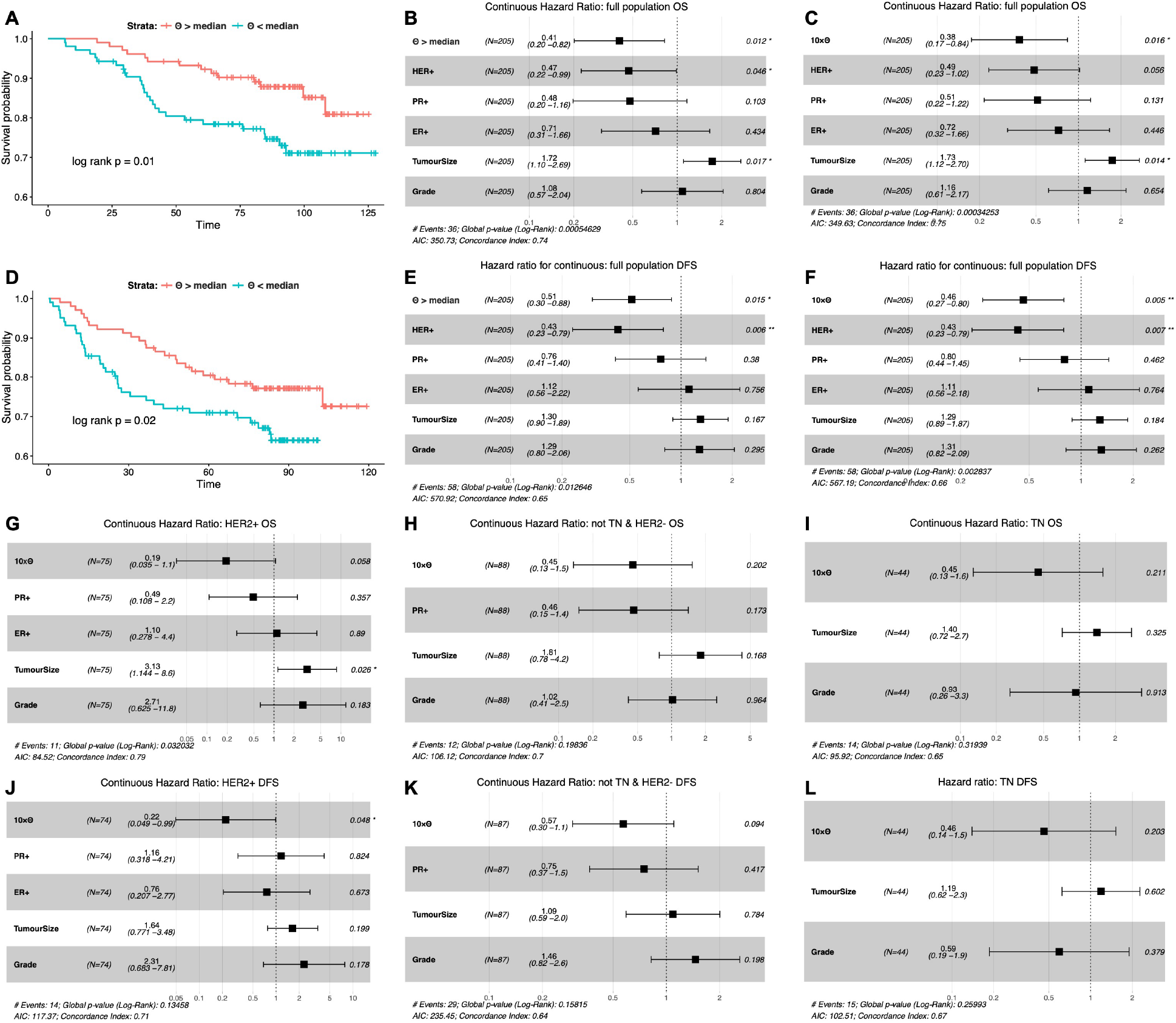
Survival analysis. (A,D) Kaplan-Meier survival plot showing differences in overall survival (OS) and disease-free survival (DFS) for patients above and below the median Θ value. (B,E) Multivariate analysis of overall survival using the Cox Proportional Hazards Model to calculate hazard ratios (HRs) for the patients above the median Θ value compared to those below it. Only those with Θ < 0.3 are included because those with Θ ≥ 0.3 have likelihood curves with significant flat contributions to Θ (Note S2.2, Fig. S2). (C,F – L) Multivariate analysis of OS or DFS using the Cox Proportional Hazards Model to calculate hazard ratios (HRs) for Θ as a continuous variable (HRs for 10Θ are shown to get sizes comparable to other factors). (C,F) Whole population with Θ < 0.3. (G,J) HER2 positive patients. (H,K) Patients who are HER2- and not Triple Negative. (I,L) TN patients. See Table S3 for other prognostic strata.

This analysis confirmed that clock function was an independent prognostic factor for both OS (HR^1/10^ = 0.38 [0.17-0.84], p = 0.016) and DFS (HR^1/10^ = 0.46 [0.27-0.80], p = 0.005) (Figure 4 C, F). The worse the tumour clock dysfunction, the larger the prolongation of both OS and DFS. The other significant prognostic factor for OS was tumour size (T2 vs T3-4; p = 0.014), with a trend for HER2 status (positive vs negative; p=0.056). Strikingly, the only statistically significant prognostic factor for both OS and DFS was clock dysfunction, with tumour size also significant for OS and HER2 status for DFS. Their effects were comparable in magnitude.

Breast cancers represent a set of highly heterogeneous diseases which are usually classified according to oestrogen receptor (ER), progesterone receptor (PR) and HER2 expression status (*52, 53*). Patients with ER+ tumours represent the majority of the patients, with a good prognosis despite a poor response to chemotherapy. Patients with HER2 positive cancer have largely benefitted from anti-HER2 treatments, which have drastically improved their prognosis. Patients with triple-negative (TN) breast cancers, phenotypically defined by the lack of expression of ER and PR and that of HER2 overexpression and amplification, are both very sensitive to chemotherapy but also have the worst prognosis among the breast cancer subsets. We therefore further examine the relevance of Θ in these three categories for OS (Figure 4 G-H) and for DFS (Figure 4 J-L). We see that for the HER2+ subset, the effect on HR is very substantial with HR^1/10^ values around 0.2 for both OS and DFS. The HR^1/10^ decreases to 0.45 for OS and 0.57 for DFS in the HER2-/NonTN subset, and to 0.45 and 0.46 respectively for the TN group, suggesting a possibly lesser prognostic effect of clock dysfunction there.

For many prognostic strata the hypothesis HR < 1 is statistically significant. For OS these are the whole population, PR-, pCR-, grade 3, nodal status 1 and not TN (Table S3). For DFS this is true for the whole population, ER-, PR-, HER2-, PCR-, grade 3, nodal status 1, no LVI and not TN (Table S3). Pathological complete response (pCR) is a prognostic factor defined as absence of residual invasive cancer cells in the breast and axillary lymph nodes. It is therefore associated with tumours that display highest susceptibility to chemotherapy, and it is consequently of interest that, apart from tumour size, this established prognostic factor is the only one that is significantly correlated with Θ (Fig. S5I).

The above results raise the question of whether correlations with other known explanatory factors fully or partially explain the link of Θ to survival. We naturally asked whether these results could be due to correlation of Θ with known prognostic factors. Of the established prognostic factors only two showed a significant relationship with Θ (Fig. S5). Those tumours that achieved pCR have a higher median Θ than those that did not, as did those large tumours (T3-T4) compared with the smaller ones (T2) (Fig. S5I). Moreover, the multivariate analyses using the Cox model provides strong evidence of independence of Θ from the classical prognostic factors, as the hazard ratio HR and the p-value for Θ are much the same for the univariate and multivariate analyses. If Θ was correlated with any of these factors, we would expect to see an HR value closer to 1 when the multivariate analysis is compared to the univariate one. Using a multivariate Cox analysis, Cadenas *et al.* (*40*) showed that of the clock-related genes only *PER3*, *RORC* and *TIMELESS* were associated with survival independently of other prognostic factors. All four *PER3* probes have a Spearman correlation with Θ not significantly different from 0 and, of the two probes of *RORC*, one is significant with a negative Spearman correlation of −0.2. However, since overexpression of *RORC* is associated with longer MFS, it acts in the same way to Θ and therefore the negative correlation implies *RORC*’s association with survival is unrelated to that of Θ. *TIMELESS* is more relevant because probe “203046_s_at” has a relatively large negative effect on the HR and a p-value < 0.001. However, its correlation coefficient with Θ is −0.0353 (p = 0.5977) implying that it also does not explain the significance of Θ. We also tested for correlation of Θ with the PCNA meta-gene set that is consistently associated with cell proliferation and found that any correlation is weak or non-existent (Note S10.5). Taken together, these analyses reveal that the circadian clock function of tumours, measured by our metric Θ, adds significant information independently of the other prognostic factors and represents itself a potentially useful prognostic and predictive biomarker. Its effect on hazard is substantial and on a par with the most significant prognostic factors.

### TimeTeller likelihood curves have the flexibility to assess functionality

As discussed above and in Note S4 the variance of the sample timing MLE *T* depends crucially on the variance structure of *P*(*g*|*t*). The above analysis of the TimeTeller likelihood curves shows them to be highly flexible and able to contain interesting biological information. While the ZeitZeiger likelihood curves ((*24*), Note S13.2), may be more accurate at assessing timing they are much more uniform (Fig. S5G) and do not contain this information (Note S10.4, Fig. S5E,F) presumably because they are not transparently connected to *P*(*g*|*t*). They have very uniform Θs across the REMAGUS data and are only very weakly correlated with the TimeTeller Θs (Fig. S5).

While the likelihood curve associated with samples from the training data have a single peak, many samples with poor clock function can have more than one (Figs. S2). Indeed, a characteristic of likelihood curves corresponding to poor clock function is the emergence of a second peak more than 6h from the dominant peak as shown in Figs. S2. The reason for this is that for some times *t* the training data points at CT*t* are close to those of CT(*t* + *t*_1_) with *t*_1_ ≈ 8h – 16h (Fig. S5) and this can give rise to two peaks in the likelihood curve *L*_*g*_(*t*) if the data point corresponding to *g* lies roughly equidistant from the training data points at CT*t* and those at CT(*t* + *t*_1_). In samples with poor clock function we often find that the second peak gives a reasonable time estimate when the primary peak time is unreasonable (Figs. S4D & S5(A-D,G)). For example, in the Feng et al. and REMAGUS cancer datasets a large proportion of the samples with clearly wrong timing using the primary peak moves to a timing within the timing interval defined by those REMAGUS samples for which we have times (Figs. S4D & S5A,B).

This raises the question of which aspects of the likelihood curves can identify interesting aspects of disease and, in particular, to what extent the presence/absence of a second peak is what is determining the significant hazard ratios found in the REMAGUS data. Analysis (Note S10.6) of this shows that even if we restrict attention to those samples with a single peak (so that Θ corresponds to the classical statistical construct for confidence intervals) or to only samples giving multiple peaks (where our extended definition of Θ applies) there is still good association with survival strongly suggesting that both the classical construct and the extended one used here provide interesting biological information.

## Discussion

Our study had two aims. Firstly, to provide a way of assessing from a single biological sample how well the circadian clock is working, and secondly, to highlight its relevance for circadian medicine through a stringent test. We applied TimeTeller to breast cancer transcriptome data and show that survival and our definition of clock functionality were strongly linked.

### Assessing clock dysfunction

TimeTeller produces an estimate Θ of clock functionality that is based on the likelihood curve *L*_*g*_(*t*) that it estimates. The key to why this works well is the correlation structure in the data points at a given time. The *G* clock genes in our training data are far from independent and although they are noisy and subject to measurement error, they have a clear correlation structure and their covariance matrix has rapidly decreasing eigenvalues. This means that, considered as a vector, they can have an accuracy in assessing time *T* that is much greater than any single gene. Thus, we believe that our multi-dimensional approach studying the data in G-dimensional space and combining several dimension-reducing projections is crucial.

Although TimeTeller does a reasonable job at assessing timing we believe that ZeitZeiger is currently more accurate for this. For example, for the REMAGUS data ZeitZeiger estimates the time range of the data better and does not have to rely on second-peak corrections. On the other hand, TimeTeller does a reasonable job on timing and, unlike other algorithms, provides functionality scores from individual samples (S11.4). Clearly future work should involve combining the best aspects of each of these algorithms.

In our discussion here we have restricted attention to the circadian clock but there is no reason in principle why this approach cannot be applied much more generally. For example, it would be of great interest to apply it to a coupled system such as that involving the circadian clock and cell cycle, or to the clock and any representative set of rhythmic downstream genes. Indeed, it is worth noting that one of the genes identified in the mouse model is the gene *Wee1* which provides a key connection between the clock and cell cycle (*54*).

Further work is needed to try and understand what aspects of the cells and tissues give rise to the high Θ values we observe in diseased tissue. Since our metric Θ gives a stratification of clock function in cells and tissue we can use this to define a more targeted search to uncover the links between the clock and the mechanisms leading to disease and cellular dysfunction.

The key outputs of our algorithm are the likelihood curve and the associated LRF and the metric Θ is just one summary statistic. There may well be other summary statistics that will be of use for characterizing the likelihood curve and we feel that when considering a single sample one should always inspect the likelihood curve and LRF to assess the quality of the analysis. The maximum value of the likelihood curve (i.e. *P*(*g*|*T*)) is often very informative. Moreover, in clinical use one would know the timing and this would allow one to extract more information from the likelihood function, for example to assess if a second peak gives the correct timimg.

### Breast cancer survival

We have shown very clear and consistent links between tumour Θ values and both 10-year survival and 10-year disease-free survival in the breast cancer patients from the REMAGUS trial. Despite the large body of work showing that disruption of the host clocks was associated with poor prognosis and that chemotherapy timing could make the difference between life or death in preclinical breast cancer (*19–22*), there was previously no simple method which would allow its measurement in daily oncology practice. Our work has the potential to change this as the method we present only requires a single sample and can be adapted to any gene expression technology. We envisage the use of this metric in conjunction with current prognostic factors to refine treatment management. The results should also open up new opportunities for research into the circadian clock as a target for treatment using the stratification by the dysfunction metric. The techniques developed here can potentially be applied to other diseases involving the circadian clock and to other regulatory systems by extending the gene panel outside of the circadian clock as discussed above.

Since the effect on the hazard rate is so substantial it is important to try and understand the mechanisms behind this. In considering our results it is important to distinguish the functionality of the host clock from that of tumour cells. Our analysis suggests a significant survival advantage when the tumour clock is dysfunctional and one is tempted to conjecture that the contrast between this and good host rhythmicity is behind this observation. It has previously been observed that severe circadian clock disruption in experimental cancer cells caused by *Cry1*/*Cry2* double knock out improved the efficacy of chemotherapy (*55*). This, together with our observation of correlation between high Θ and pCR hints at a link with increased sensitivity to chemotherapy. As mentioned above, the stratification of tumours by Θ might provide an experimental strategy for understanding this. Overall, we have developed a new model for tissue clock functionality that could help refine treatment strategies for breast cancer. We expect that the use of algorithms such as TimeTeller will contribute to circadian medicine at large by enabling the integration of molecular clock determinations in diseased tissues with the design of innovative clock-targeted therapies.

## Code and data availability

The code and data underlying the results of this study are available for reviewers and will be made publicly available on publication.

## Acknowledgements

We thank Ida Iurisci (INSERM, Villejuif) and Anne-Sophie Hamy (Paris Descartes University) for help with complementary information regarding the breast cancer database, and Meritxell Saez and Laura Usselmann (University of Warwick) for a critical reading. DAR and DV thank the Engineering & Physical Sciences Research Council (EPSRC) for a PhD studentship through the MOAC Doctoral Training Centre grant number EP/F500378/1. DAR was also supported by Biotechnology and Biological Sciences Research Council (BBSRC) Grant BB/K003097/1 and EPSRC Grant EP/P019811/1. FL & DAR were supported by a grant from Cancer Research UK and EPSRC (C53561/A19933). GAB was supported by The Anna-Liisa Farquharson Chair in Renal Cell Cancer Research.

## Supplementary Material for the paper

### S1 Probability Model & Likelihood curves *L*_*g*_(*t*)

An outline pseudocode description of the algorithm is in Note S3.

#### S1.1 Probability model

For each observation *j* = 1, …, *N* and each time where *i* = 1, … , *s*, the training data for each set of *G* expression levels is stored in the expression profiles which are *G*-dimensional vectors 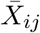. The observations *j* correspond to tissues for the mouse data and individuals for the human data. Each 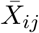 is then normalised to have a mean of 0 and standard deviation of 1, resulting in the vector *X*_*ij*_. These are the vectors that will be used to parameterise TimeTeller. Batch effects are not a problem for the data considered here as is explained in Note S4.4.

To construct the probability model we firstly construct one for each timepoint by using the local statistical structure of the data at that timepoint and then we combine these. Fix a timepoint *t*_*i*_. Associated with this is the set of *N* points *X*_*ij*_, *j* = 1, …, *N* in *G*-dimensional space. The projection operator described in Note S2 gives an optimal way to linearly project these points into *d*-dimensional space for all *d* < *G*. This produces a corresponding set of *N d*-dimensional points *Q*_*i,j*_, *j* = 1, … , *N*. We then fit a multivariate normal distribution (MVN) to the points *Q*_*i,j*_. The dimensionality *d* is chosen so that there are enough vectors *Q*_*i,j*_ to fit a *d*-dimensional multivariate Gaussian (using the MATLAB function fitgmdist) while ensuring that most of the variance in the data is captured by the *d*-dimensional projection. In our case we take *d* = 3. A MVN distribution is defined by its mean and covariance matrix which we denote by *μ*_*i*_(*t*_*i*_) and Σ_*i*_(*t*_*i*_) respectively.

We fit a shape-preserving smoothing cubic periodic spline through the mean vectors *μ*_*i*_(*t*_*i*_) and each of the six entries that determine the 3 × 3 symmetric matrix Σ_*i*_(*t*_*i*_) so as to extend *μ*_*i*_(*t*_*i*_) and Σ_*i*_(*t*_*i*_) to all times *t* between the time points, thus obtaining *μ*_*i*_(*t*) and Σ_*i*_(*t*). A piecewise cubic Hermite interpolating polynomial spline is used in this case. This type of spline is shape preserving, i.e. continuity of the second derivative is not obligatory. This is suitable as, for example, if two covariance matrix entries were identical for two consecutive time Gaussians, the Hermite spline allows the value of the joining spline to stay the same in the space between, while a standard spline would enforce some change. This spline also interpolates so that it passes through all points. The calculations were carried out using the MATLAB function *pchip*. Using this approach, for this value of *i*, we have determined a family of MVN distributions for all times *t* between the first and last data times.

#### S1.2 *L*_*g*_(*t*) and the log threshold *l*_thresh_

Now we define the likelihood curve *L*_*g*_(*t*) where *g* is an expression profile and X is the *G*-dimensional vector obtained from *g* after normalisation. Using the idea that we define the likelihood associated with the *i*th timepoint using the probability given by the MVN i.e.

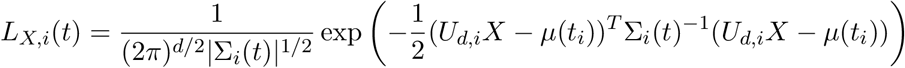

and then *L*_*g*_(*t*) is given by combining the individual likelihoods *L*_*g,i*_, *i* = 1, …, *s* as follows

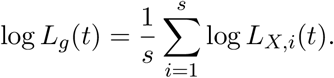

However, to ensure that this product is not wrecked by inaccurate exceptionally low values of one *L*_*X,i*_ affecting robust high values of another at the same *t* we truncate the *L*_*X,i*_. A curve *L*_*X,i*_ will take on very low values away from its maximum and these may well be inaccurate. If this happens at a *t* value for which another such curve *L*_*X,j*_ has a high accurate value then this may badly affect the estimate of *L*_*g*_(*t*). To overcome this we fix a lower threshold *l*_thresh_ < 0 and replace each log *L*_*X,i*_ by max{log *L*_*g,i*_, *l*_thresh_} in the above product.

**Figure S1:**
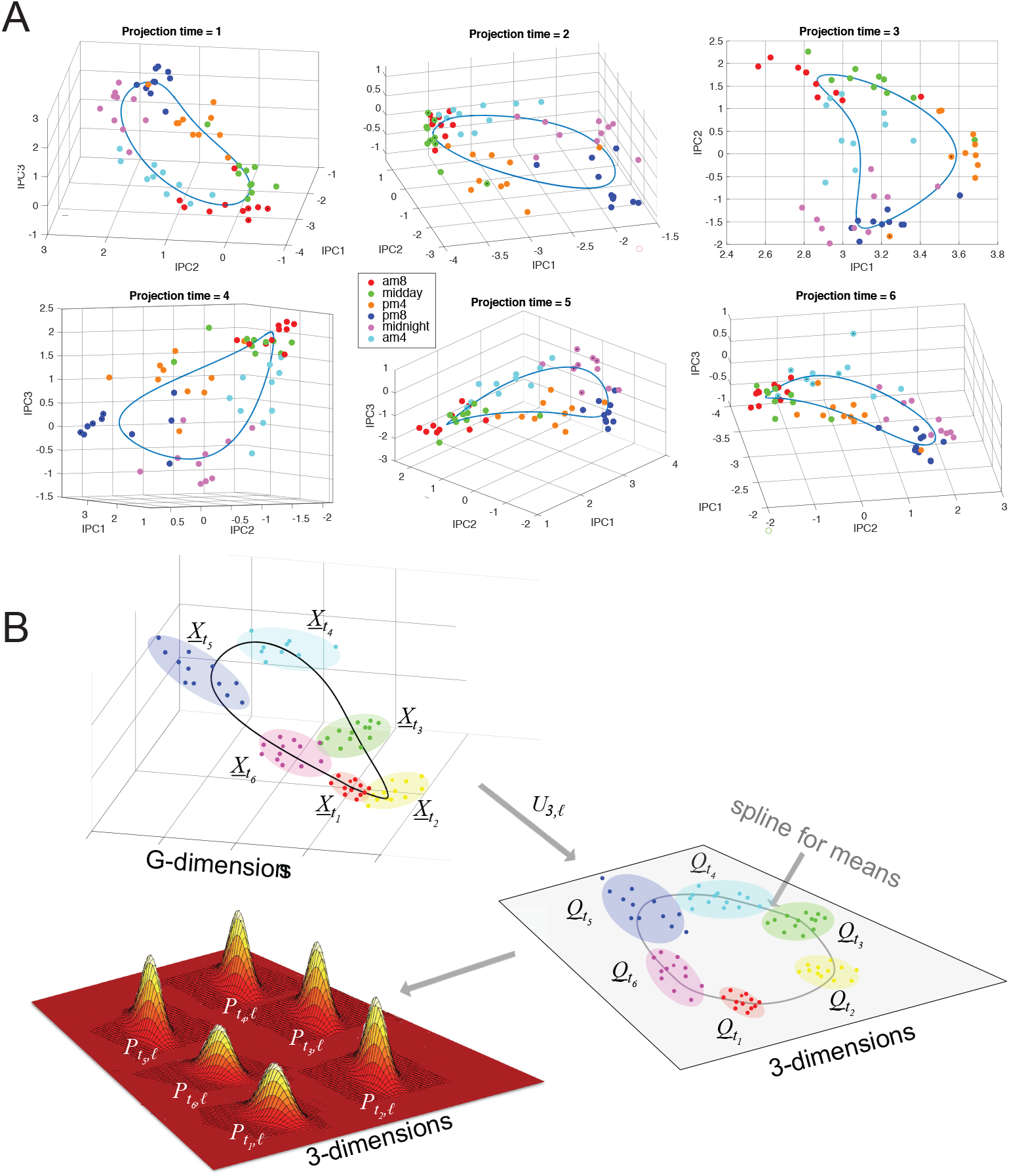
**A. All 6 local projected spaces for human data.** The data is projected as described in Material and Methods using the projections *U*_*d,i*_ where *d* = 3 and *i* indexes the six times. Each time point is coloured according to its time. Splines through the means *μ*_*i*_(*t*_*j*_) of each set of 10 time data points show (distorted) elliptical shapes. Because of this shape we often have the situation where training data points at CT*t* are close to those of CT(*t* + 12) and this can give rise to two peaks in the likelihood curve *L*_*X*_ (*t*) if the data point corresponding to *X* lies between the training data points at CT*t* and those of CT(*t*+12). **B. This schematic outlines the construction of the likelihood** *L*_*X,i*_(*t*). For each *t*_*i*_ the set of vectors 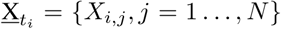 are projected into *d* = 3 dimensions using *U*_*d,i*_ to get 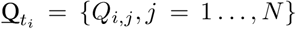. A MVN distribution is estimated for each 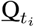 and then these distributions are interpolated using splines to all times *t* of the day. The projections *U*_*d,i*_ for the Bjarnason *et al.* data is shown above in A.

As a result of this modification, if the maximum value of log *L*_*g*_(*t*) is only just above *l*_thresh_, then it and the corresponding likelihood ratio curve will have intervals on which they are flat. This is, for example, the case for some of the curves in Fig. S2 below.

If this is the case then the length of these flat intervals above the curve *C*(*t*|*T*) defined below in Note S2 contributes to Θ. This contribution has interesting information in it because it is related to how low the maximum value of *L*_*g*_(*t*) is. However, since the size of this contribution depends upon the particular choice of *l*_thresh_ we wish to avoid this and so we generally choose a value of *l*_thresh_ that for the datasets under consideration keeps the numbers of samples with this issue as small as possible. As a result for the human data we take *l*_thresh_ = −12 or −10 and for the mouse data we generally take *l*_thresh_ = −5 or −7.

### S2 Definition of Θ

A characteristic of likelihood curves corresponding to poor clock function is the emergence of a second peak roughly 10-12h from the dominant peak as shown in Fig. S2(D,E). This shows the likelihood curves and centered likelihood ratio curves for the REMGAGUS samples with the highest second peak. The reason for this can be seen in Fig. S1(A). The curve in *d*-dimensional space tracing out the means of *P* (*g*|*t*) has a (distorted) elliptical shapes. Because of this shape we often have the situation where training data points at CT*t* are close to those of CT(*t* + *t*_1_) with *t*_1_ ≈ 8*h* 16*h* and this can give rise to two peaks in the likelihood curve *L*_*g*_(*t*).

Another way this affects our analysis is shown in Figs. S4 and S5. We see there that if we plot the real times against estimated times for those samples in the REMAGUS trial for which the sample time was recorded the data groups into two clusters, one with apparently reasonable times and the other with unreasonable times. These two clusters are roughly 12 hours apart and this suggests that for the samples that are apparently incorrectly timed the algorithm has found the wrong peak as the maximal one. Indeed, we found the second peak for all the incorrectly timed examples and found that, for the great majority, this give a seemingly correct time.

If there is such a second peak then we want to penalise it if it has a height whose ratio to the height of the primary peak is above a threshold that depends upon the time δ*t* between these two peaks. Moreover, this threshold should be minimal when δ*t* = 12h. To do this we use the curve *C*(*t*|*T*) below.

The *clock dysfunction metric* Θ is defined to be the proportion of time *t* which satisfies

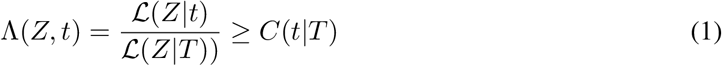

i. e. the proportion of time in the day when the given likelihood ratio curve is above the curve *C*(*t*|*T*). Here *C*(*t*|*T*) is defined by

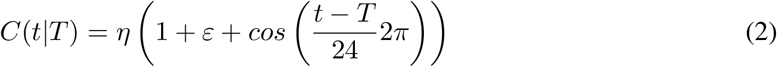

e.g. the black curve in Fig. S2. This is a simple cosine curve transformed so that

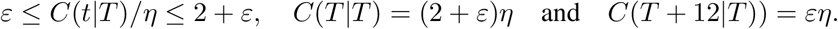

#### S2.1 Parameters *ε* and *η*

The values of *ε* and *η* used in the definition of Θ must be appropriately chosen. We must choose *ε* > 0 so that *C* > 0. The larger *ε* is, the less anti-phase peaks impact the final confidence metric.

From (1), it is clear that to ensure the positivity of Θ we must enforce that

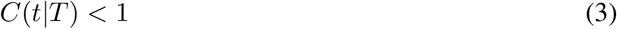

As max(*C*) = *η*(2 + *ε*) and min(*C*) = *ηε*, we must have,

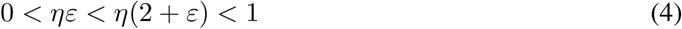

and, note that the last part of this implies 0 < *η* < 1/(2 + *ε*) < 0.5.

Also note that *ηε* defines the minimum value of the threshold for which the graph of the likelihood ratio Λ can be intersected, and *η*(2 + *ε*) defines the maximum. We set *ε* = 0.4 and *η* = 0.35, which sets the maximum threshold value (at the MLE) to *η*(2 + *ε*) = 0.84, and the minimum threshold (anti-phase to the MLE) to *ηε* = 0.14. This means that if there is a peak in the likelihood exactly anti-phase to the MLE that is more than 14% of the maximum, then it penalises Θ.

The hyperparameters *ε* = 0.4 and *η* = 0.35 are used throughout. However, we note that, since the likelihood curves are continuous (and smooth away from the ends of flat regions) the same is true of the dependence of Θ upon the parameters *η* and *ε*. Consequently, the precise choice of *ε* and *η* is not crucial.

As *ε* → ∞ the method is the same as a non penalised approach and as *ε* → 0 any anti-phase secondary peak above *m*_*t*_ will penalise Θ. An example of this is shown in in Fig. ??.

#### S2.2 Examples showing likelihood ratio curves, the curve *C*(*t*|*T*) and the corresponding Θ

In Fig. S2 we show a selection of centered likelihood ratio curves for the indicated data sets together with the graph of *C*(*t*|*T*) in black. In each case the likelihood ratio curve has been centered so that the maximum is at *t* = 12. This enables easier comparison of the curves and the associated Θ value.

**Figure S2:**
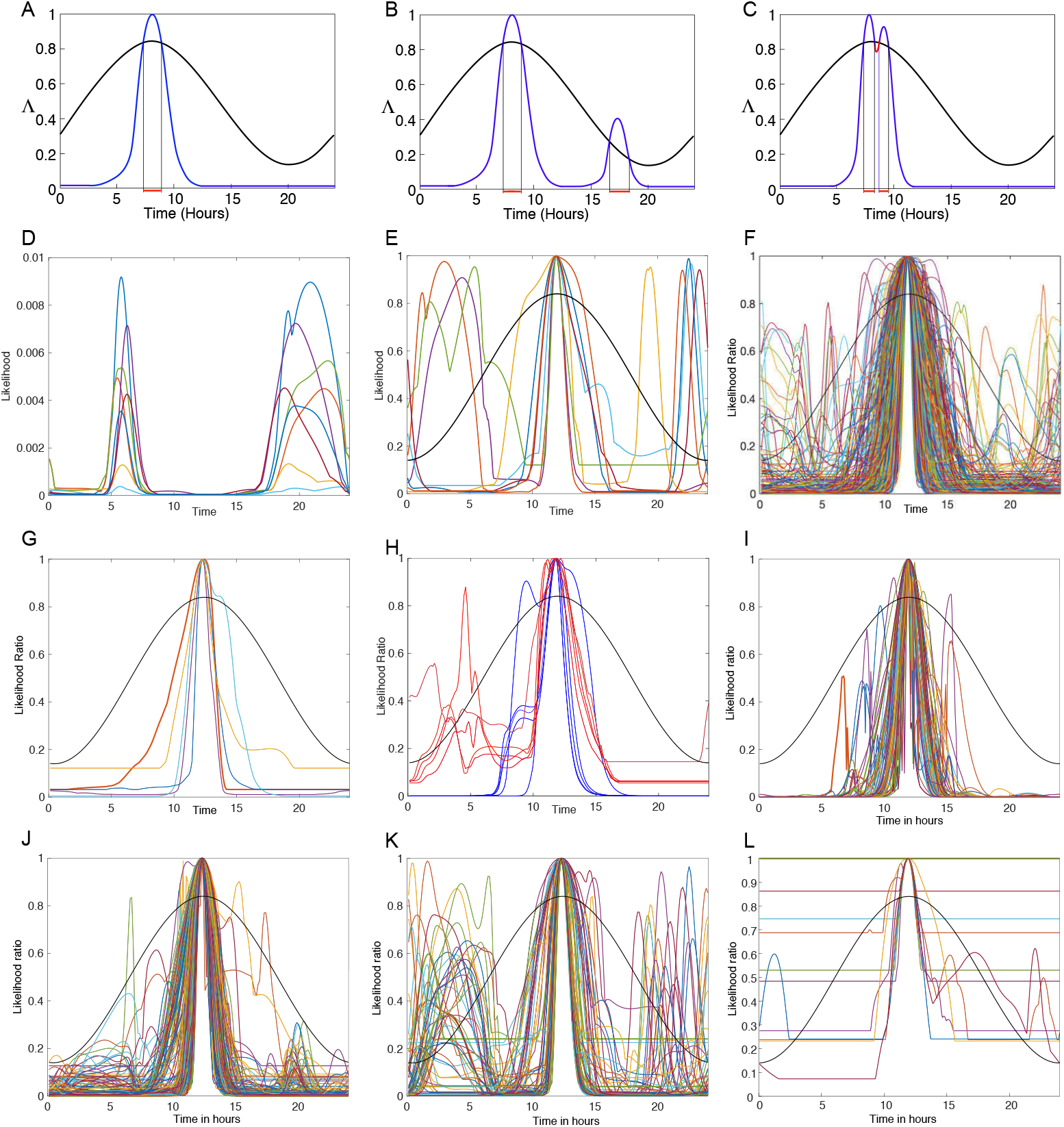
**A-C. Showing how the modification *C*(*t*|*T*) affects the definition of Θ.** The graph of *C*(*t*|*T*) is shown in black. (A) This shows how a single peak in a likelihood curve would not be affected by the penalised cut-off. (C) This shows how if there is a secondary peak near to the MLE peak, then Θ will not be penalised by much. (B) The middle panel shows how if there is a secondary peak approximately 12h from the MLE, then it will penalise Θ. The values of Θ in these plots is *l*/24 where *l* is the sum of the lengths in hours of the red segments at the base of the graph. To summarise, we want our metric to have the following properties: (i) a single, narrow, high, peak would result in a high confidence estimate; (ii) a wider peak would result in a comparatively lower confidence estimate; (iii) a second slightly lower peak near to the MLE would result in good confidence in the MLE; (iv) a low second peak around 12 hours would result in good confidence in the MLE; (vi) a higher second peak around 12 hours would result in lower confidence in the MLE; and (vii) a uniformly flat likelihood function, or one with low peak(s), would provide a low confidence estimate. **D,E. Examples of likelihood curves with two peaks.** D. These are from the samples from the REMAGUS clinical trial with the highest secondary peaks. E. The corresponding centered likelihood ratio curves (LRCs). The curve has been centered so that the maximum is at *t* = 12 to enable easier comparison of the curves and the associated Θ value. **F-H. Examples of centered likelihood ratio (LR) curves from the mouse.** In these *l*_thresh_ = -−7. F. Examples from the Zhang *et al.* data. About 15% of the curves have a second peak that intersects *C*(*t*|*T*). G. The LR curves from the Martelot *et al.* data. H. The LR curves from the Fang *et al.* data. The blue curves are for the data from the WT samples and the red curves from the KO samples. **I-L. LR curves for human data.** Here *l*_thresh_ = − 12. I. The LR curves from the Bjarnason human data. J-L. The curves for the REMAGUS data with Θ < 0.1 (J),0.1 < Θ < 0.3 (**K**) and 0.3 < Θ (L). For the last dataset the flat part of the LR curves intersect the *C*(*t*|*T*) curve (black) and therefore the Θ values are significantly affected by this. This occurs because the maximum value of the corresponding likelihood curve is close to the threshold value *l*_thresh_ used. To overcome this one can decrease *l*_thresh_.

### S3 Outline pseudocode of algorithm

**Algorithm 1:**
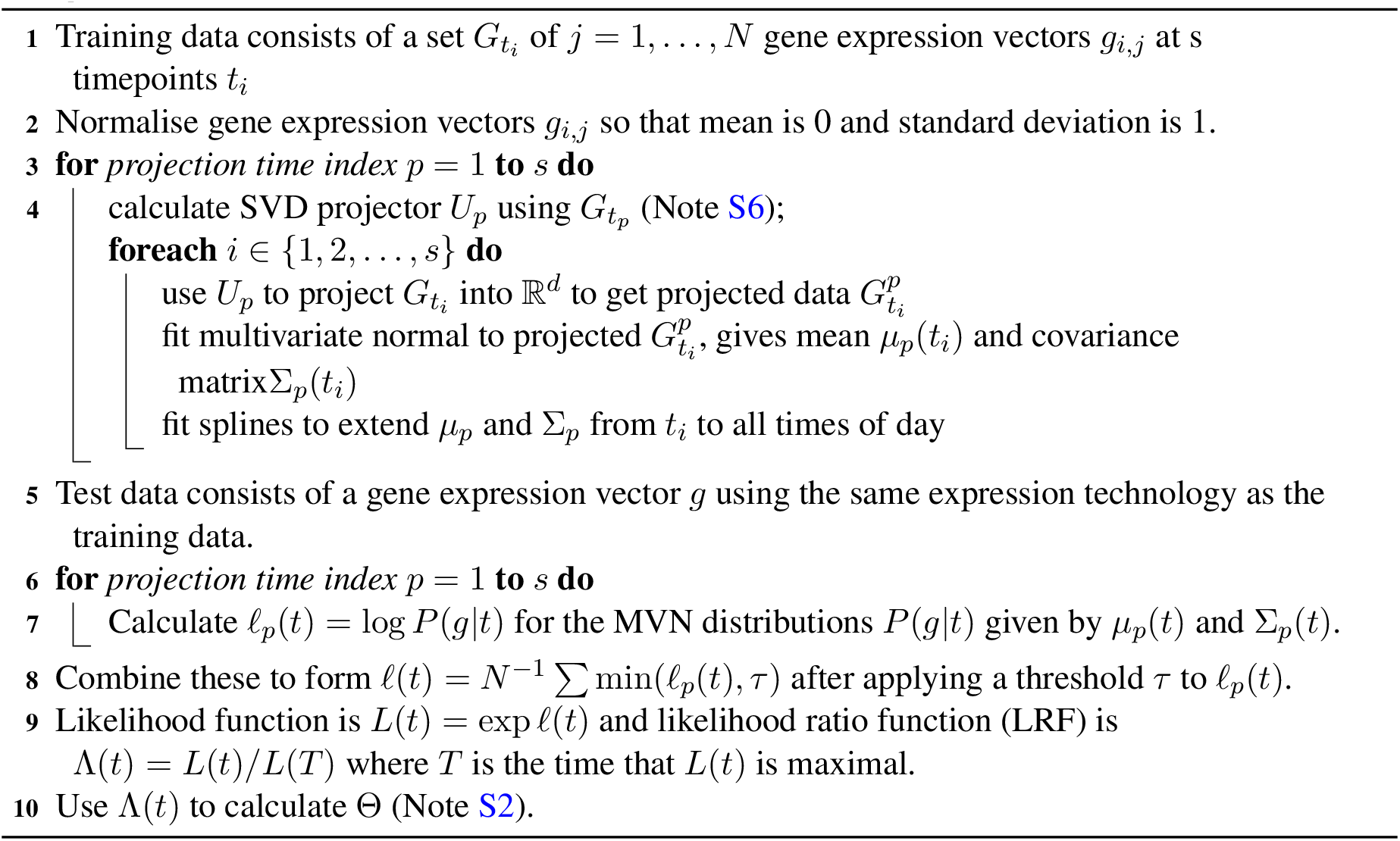
TimeTeller.

### S4 Role of gene-to-gene correlations in the likelihood function

#### S4.1 Θ contains an estimate of a direct measure of clock precision

In what follows *g* = (*g*_1_, …, *g*_*G*_) denotes a vector consisting of the possibly normalised levels of *G* clock-associated genes. Call these vectors rhythmic expression profiles.

We consider the probability *P* (*t, g*) that *g* is observed at time *t* in WT conditions and the associated conditional probability distributions *P* (*g*|*t*) and *P* (*t*|*g*). Then if we want to assess how well the clock is working in an independent sample with expression vector *g*^*^ ℝ^*G*^ we would want to estimate the conditional distribution *P* (*t*|*g*^*^). The time *T* would be estimated to be the value of *t* maximising *P* (*t*|*g*^*^) i.e. the maximum likelihood estimate (MLE).

The corresponding likelihood ratio is given by

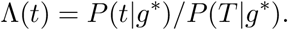

As is well-known (e.g. Chaps. 8 & 9 of (1)) the likelihood ratio confidence interval gives a good estimate of the confidence intervals for the MLE when the log likelihood function is approximately quadratic on some scale as is the case with our likelihood functions. If α is the required sensitivity (i.e. 1 minus the confidence level) then the confidence interval is given by the set of times *t* that satisfy

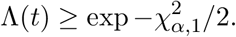

We can relate this to Θ (see Note S2) when the likelihood curve only has a single peak (or, more generally, only twice meets the curve *C* in Note S2). Since the likelihood ratio curve is quadratic near its maximum we deduce that the contribution to Θ from the region around the maximum is

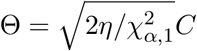

where *C* is the length of the confidence interval and *η* is the parameter in Note S2.1. Since the confidence intervals define the precision of the clock we see that, in this case, Θ is a direct measure of clock precision.

#### S4.2 P (t|g) is estimated using *P*(*g*|*t*)

To measure *P* (*t*|*g*) we will use the fact that, by Bayes theorem, so far as dependence upon *t* is concerned,

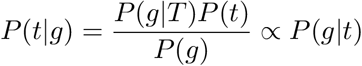

provided we assume that *P* (*t*) is reasonably independent of *t*. Therefore, we can use *P* (*g*|*t*)/*P* (*g*|*T*) to estimate Λ(*t*) = *P* (*t*|*g*)/*P* (*T* |*g*). Consequently, TimeTeller tries to estimate *P* (*g*|*t*) directly from WT data. For the estimation it is assumed that *P* (*g*|*t*) is multivariate normal.

#### S4.3 Θ depends crucially on the covariance structure of *P*(*g*|*t*)

To see what Θ depends upon consider the case where *P* (*t*|*g*^*^) is relatively sharply peaked around *T*. Expanding around this estimate, the distribution is approximately Gaussian with

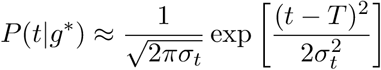

and the variance *σ*_*t*_ is given by

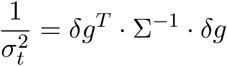

where Σ is the covariance matrix of *P* (*g*|*t*) and *δ*_*g*_ is the derivative with respect to *t* of the mean of *P* (*g*|*t*) at *g* = *g*^*^*, t* = *T*. Indeed, if we drop the tightness assumption we can use the Kramer-Rao theorem to deduce that the term on the righthand side is a lower bound for the variance because this term is the dominant part of the Fisher Information matrix of *P* (*g*|*t*) with respect to *t*.

Therefore, we see that it is crucial in estimating our clock dysfunction metric that we take account of the covariance structure of *P* (*g*|*t*).

### S5 Data sets

#### S5.1 Training datasets

##### Mouse: Zhang *et al.* mouse timecourse microarray data

Zhang *et al.* (*2*) quantified the transcriptomes of 12 mouse tissues every 2 hours over 48 hours with microarrays, and every 6 hours using RNA-seq. The tissues sampled were from the adrenal gland, aorta, brainstem, brown fat, cerebellum, heart, hypothalamus, kidney, liver, lung, skeletal tissue, and white fat of the mice. Four of the tissues were excluded from our analysis (3 brain tissues and white fat) as the noise to amplitude ratios are higher for these.

###### Experimental setup

6 wk-old male C57/BL6 mice were entrained to a 12h:12h LD schedule for 1 week, and then released into constant darkness (CT0). The mice were provided with food and water *ad libitum*. From CT18 post-release, 3 mice were killed every 2 hours, for 48 hours (until CT64), and specimens from 12 organs were snap frozen in liquid nitrogen.

###### Microarray methods

RNA was extracted and pooled for three mice for each tissue and time point. RNA abundances were quantified using Affymetrix MoGene 1.0 ST arrays using the standard manufacturer’s protocol. The results of this make up the .CEL files that are used in this study. They are accessible via GEO accession number GSE54652.

In their analysis, Zhang *et al.* used the RMA algorithm to normalise the data using the Affymetrix Expression Console Software. As is standard, GeneChips were matched to genes and filtered for protein-coding status, resulting in 19,788 genes forming the background set. They performed oscillation detection using the R package JTK CYCLE (*3*) to fit time series data and detect oscillations on transcripts, using a the cutoff significance of q<0.05 (5% FDR).

##### Human: Oral Mucosa Timecourse microarray data (Bjarnason *et al.*)

For this study Bjarnason *et al.* (*4*) recruited ten healthy human volunteers, five female and five male. Mucosa tissue was collected at six time points: 8 am, noon, 4 pm, 8 pm, 12 midnight, and 4 am. Subjects were selected after screening by clinical history, physical examination, routine blood work (complete blood count, electrolytes, creatinine) and actigraphy to confirm regular sleep-wake patterns. Mucosa samples were collected by a dental surgeon, using a tissue punch biopsy. Subjects went to sleep in a darkroom at their usual bedtime and were awoken for the midnight and the 4 am samples^1^.

After tissue mucosa samples were collected, they were immediately frozen in liquid nitrogen and stored at −80°*C* until use. Total RNA was prepared by Trizol Reagent (Invitrogen) in accordance with the manufacturer’s specifications. RNA samples were quantified by optical density measurements at A260nm and A280nm. All samples were determined to be of high quality with A260:A280 ratios > 1.9. Total RNA (5*μ*g) of each sample was used for microarray analysis on Affymetrix HG U133 Plus2 chips. Cumulatively this chip represents 54,679 gene transcripts for analysis. Biotinylated cRNA was prepared according to the standard Affymetrix protocol (Expression Analysis Technical Manual, 2004, Affymetrix). Following fragmentation, 15 *μ*g of cRNA were hybridized for 16 hrs at 45°*C* on GeneChip Human Genome U133 Plus 2.0 arrays. GeneChips were washed and stained in the Affymetrix Fluidics Station 450 and were scanned using the Affymetrix GeneChip Scanner 3000.

Although there were 16 probes identified in Note S7 as rhythmic and synchronised, we only use 15 probes going forward. The reason is that the Per1 probe 244677 at was found to have significant signal issues in many of the independent human datasets, i.e. the signals values were very low. As there is another Per1 probe in this dataset that does not have this problem, we can conclude that this is a probe issue, and not an issue with the Per1 gene expression.

#### S5.2 Other mouse datasets

We also consider the following datasets which are comparable because (i) they use the same male mouse C57BL/6 and (ii) all microarray expression analysis used the Affymetrix MoGene 1.0 ST GeneChip.

- LeMartelot *et al.* (*5*). This is made up of microarrays from pooled RNA extracted from the whole liver of 5 mice. Samples were collected at times ZT2, ZT6, ZT10, ZT14, ZT18, ZT22, ZT2(+24) after the mice had been entrained to LD cycles for 2 weeks. Accessible via GEO accession number GSE35789.
- Fang *et al.* (*6*). 5 Wild Type and 5 Rev-erbα KO mice were entrained to LD cycles and euthanised at ZT10, where liver samples were taken. As the training data is labelled between CT18 and CT64, this corresponds to CT34 in TimeTeller. The authors report that the knock-out was successful. Accessible via GEO accession number GSE59460.
- Barclay *et al.* (*7*). All mice were WT but half of them were sleep deprived when they would usually be sleeping. All mice were synchronised to LD cycles and given food *ad libitum*. The group of stressed mice were kept awake during the first 6 hours of light (ZT 0-6) on days 1 through 5 and days 8 through 12 using a gentle handling approach designed to minimize stress effects and intervention by the experimenter. At all other times mice were left undisturbed, and the other group of mice were left completely undisturbed. Liver and adipose samples were taken from 3 mice in each of the 2 conditions at ZT1, ZT7, ZT13, and ZT19. Accessible via GEO accession number GSE33381.

#### S5.3 Other human datasets

Working with human data raises extra issues. The training set is smaller, and we are not working with genetically identical humans. All of the data we consider is from live tissues, and not *in vitro* samples as we expect this to be appropriate for meaningful comparison with diseased tissues. So far as we are aware the only 24h timeseries for biopsies from healthy human tissue using the U133 2.0 GeneChip is that of Bjarnason *et al.* (Note S5.1).

In order to attempt to keep the variance low, we start by testing TimeTeller on datasets originating from human oral mucosa, like the training data. The datasets considered are

- UK-Sri Lankan healthy data. Saeed *et al.* (*8*) conducted a study comparing UK and Sri Lankan oral squamous cell carcinomas. The control data for this study includes 5 healthy control oral mucosa samples from the UK and 3 healthy control samples from individuals in Sri Lanka. The study states that the samples were handled using identical protocols for tissue collection and processing^2^. All individuals were healthy, with low risk factors for cancers of the mouth, but there is no other information given on the individuals or their lifestyles. Accessible via GEO accession number GSE51010.
- Boyle *et al.* (*9*) studied the effects of smoking on the oral mucosal transcriptome. Forty current smokers and forty age and gender matched never-smokers underwent buccal biopsies. Eligible subjects were healthy volunteers (except for the effects of smoking in the smoker cohort). Since one sample was excluded from the study based on a quality measure we have 79 samples for analysis. Accessible via GEO accession number GSE51010.
- Feng *et al.* (*10*) conducted a comparative analysis of healthy oral mucosa transcriptome and oral squamous cell carcinoma (OSCC) transcriptome. The dataset contains 229 samples in total, 167 of OSCCs, 45 of normal oral mucosa, and 17 samples are of dysplasic oral mucosa tissue. Dysplasic tissue is abnormal tissue that could signify early signs of cancer. Accessible via GEO accession number GSE59460.
- REMAGUS clinical trial (*11*). This is described in detail in the main paper and (*11*) contains a detailed analysis of data quality. There are 226 analysed tumour samples with 10-year survival data and detailed clinical data. Accessible via GEO accession number GSE26639.

#### S5.4 Batch Effects

In high-throughput studies batch effects occur because measurements are affected by variations in experimental conditions such as laboratory conditions, reagent lots, and personnel differences. We used the R packages BatchQC, ComBat and sva to analyse batch effects.

For the training data we are only concerned with the *G* = 10 − 16 selected clock-associated genes. We inspected the embedding of the data into *G*-dimensional space and the subsequent projections into 3-dimensional space using SVD for any sign of batch effects especially with respect to timing of the samples (see Figs. S1A and S6). The most convincing evidence of an absence of batch effects is the analysis of error in Fig. 2C of the main paper and Fig. S3F which shows that for each individual, the deviation from the mean of the timing for each individual is consistent across times and lines up very closely with the phase of the genes shown in these figures. In any case we would not want to make batch correction as this would arbitrarily destroy the statistical distribution of the data which is a crucial ingredient of our algorithm.

For the test data there is no question of a batch effect affecting the determination of Θ in a test data set as each sample is treated individually i.e. it is treated individually under both vector normalisation and the initial fRMA normalisation and then the probability model is applied to it. For the REMAGUS data the batch effect question is also dealt with in reference (*11*) where there is a detailed analysis of data quality. Our analysis also found no evidence of batch effects in this data when projected into *d* = 3-dimensional space using PCA.

### S6 Singular Value Decomposition

We will use Singular Value Decomposition (SVD) in a number of places and therefore we outline it here. SVD gives a decomposition of any *m* × *k* matrix *A* into a product of the form *A* = *UDV* ^*^ where *U* is a *m* × *k* column-orthonormal matrix (*UU* ^*^ = *I*_*m*_ and *U* ^*^*U* = *I*_*k*_), *V* is a *k* × *k* orthonormal matrix and *D* = diag(*σ*_1_, …, *σ*_*k*_) is a diagonal matrix.^3^ This is the version of SVD that is often called thin SVD (see (*12*) for more details). The ordered elements *σ*_1_ ≥ · · · ≥ *σ*_*k*_ are the *singular values* of *A*. We note that the columns *V*_*j*_, *j* = 1, … *k*, of *V* form an orthonormal basis for ℝ^*k*^. If the last *n* of the singular values are zero then the last *n* columns are an orthonormal basis for the kernel of *A*. If *m* < *k* then *n* ≥ *k* − *m*.

In what follows we will refer to the columns *V*_*j*_ (resp. *U*_*j*_) of *V* (resp. *U*) as the right (resp. left) singular vectors. Note that *AV*_*j*_ = *σ*_*j*_*U*_*j*_ and 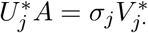.

#### S6.1 Optimal projections via SVD and the projection *U*_*d,i*_

Suppose **v** = *v*_1_, *v*_2_, …, *v*_*N*_ is a set of *N n*-dimensional data vectors. Let b = *b*_1_*, … , b*_*n*_ be an orthogonal basis of unit vectors for ℝ*n*. Given b consider the *k*-dimensional projection of a vector *v* in **v** given by

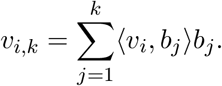

If the mean error *e*_*k*_(**b**) of this projection is defined by 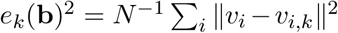 we seek a basis which minimizes this error for all *k* ≤ *n*.

Define *σ*_*i*_(**b**) by

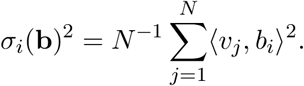

then, by orthogonality of the *b*_*i*_, 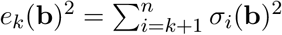.

As is well-known in (e.g. (*12*) (*13*)), the solution to this problem is provided by SVD. If *M* is the *N* × *n* matrix whose columns are the vectors *v*_*i*_, let *M* = *UDV* ^*^ be the SVD of *M* (* denoting transpose). Then *U* is *N* × *n* and the solution to the above problem is *b*_*i*_ = *U*_*i*_, the *i*th column of *U*, and the *σ*_*i*_(**b**) are the singular values of *M* (i.e. the diagonal elements of *D*). Moreover, projecting into ℝ^*k*^ using this basis gives

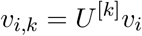

where *U* ^[*k*]^ is the transpose of the matrix whose *k* columns are *U*_1_, …, *U*_*k*_.

Thus for all *k* = 1, …, *n* the projection of the vectors into *k* dimensions using *U* ^[*k*]^ is the optimal representation of the vectors in ℝ^*k*^ in the sense that the mean *L*^2^ error is minimal.

The projection *U* ^[*d*]^ defined above is defined in terms of the vectors **v** = *v*_1_*, v*_2_, …, *v*_*N*_. If v consists of the vectors *X*_*i,j*_ for *j* = 1, … , *N* (i.e. *v*_*j*_ = *X*_*i,j*_), where *X*_*i,j*_ is as defined in Note S1 then we denote the projection *U* ^[*d*]^ by *U*_*d,i*_.

### S7 Analysis of synchronicity and rhythmicity

To measure synchronicity amongst individuals we used the approach using Singular Value Decomposition (SVD) as explained in section S6. We describe this in the context of the Zhang *et al.* data.

After fRMA processing, the data contains 35,556 probe values for *m* = 8 organs at *T* time points which was structured into 35,556 *T* × *m* matrices *M*_*g*_, and normalised so that each *T*-dimensional column has mean 0 and standard deviation 1. The columns thus correspond to normalised timeseries and we have one for each organ. Then if the SVD of these matrices has the form 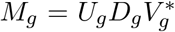 as in Note S6, the optimal projection, as in Note S6.1 is given by using as basis the columns *U*_*g,i*_ of *U*_*g*_ which correspond to timeseries.

Thus, the first principal component *U*_*g*,1_ describes the dominant temporal shape found across the various organs for gene *g* and the other principal components *U*_*g,i*_, *i* > 1, describe how this is modified across organs in a graded way with the strength given by the singular values *σ*_*g,k*_. To characterise the relative strength of the first principal component we use

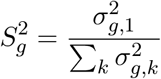

and this is our *synchronicity score*. See Table S1.

For JTK analysis the package at https://github.com/mfcovington/jtk-cycle was used. Using this we classified which genes optimise both rhythmicity and synchronicity. To choose the set of probes to make up the training set for TimeTeller a scoring of 1-50 is assigned to each probe according to its rank, and then these are simply summed up to combine results. This is shown in Table S1.

**Table S1:** Summary of results of SVD, COSINOR and JTK analysis. (top) For the Bjarnason *et al.* data. Both the SVD and COSINOR are equally weighted in a ranking to find the most rhythmic and synchronised probes. For the Zhang *et al.* data (bottom) the ranking for the JTK analysis is also added.

| Gene and ProbeID | % Var | COSINOR gmean | Ranks |  |  |  |
| --- | --- | --- | --- | --- | --- | --- |
|  |  |  | SVD | COSINOR | JTK | Sum |
| Human Data |  |  |  |  |  |  |
| PER3_221045_s_at | 0.87 | 8.00E-12 | 50 | 48 | - | 98 |
| PER3_1569701_at | 0.85 | 2.66E-13 | 46 | 50 |  | 96 |
| TEF_225840_at | 0.87 | 4.97E-11 | 49 | 47 |  | 96 |
| NR1D2_225768_at | 0.86 | 4.97E-11 | 48 | 46 |  | 94 |
| PER2_205251_at | 0.84 | 2.66E-13 | 44 | 49 |  | 93 |
| CIART_227285_at | 0.85 | 1.72E-10 | 45 | 45 |  | 90 |
| PER1_202861_at | 0.84 | 6.10E-10 | 43 | 43 |  | 86 |
| DBP_209782_s_at | 0.86 | 1.03E-08 | 47 | 39 |  | 86 |
| NR1D2_209750_at | 0.83 | 2.01E-10 | 41 | 44 |  | 85 |
| ARNTL_210971_s_at | 0.82 | 6.10E-10 | 39 | 42 |  | 81 |
| ARNTL_209824_s_at | 0.83 | 3.74E-09 | 40 | 40 |  | 80 |
| NPAS2_213462_at | 0.83 | 2.65E-07 | 42 | 36 |  | 78 |
| NR1D1_204760_s_at | 0.80 | 3.11E-09 | 36 | 41 |  | 77 |
| NR1D1_31637_s_at | 0.82 | 1.86E-08 | 38 | 38 |  | 76 |
| PER1_244677_at | 0.81 | 6.36E-08 | 37 | 37 |  | 74 |
| NPAS2_39549_at | 0.77 | 1.46E-06 | 35 | 33 |  | 68 |
| NPAS2_1557689_at | 0.73 | 1.10E-06 | 32 | 34 |  | 66 |
| HLF_204753_s_at | 0.70 | 6.49E-07 | 27 | 35 |  | 62 |
| NPAS2_1557690_x_at | 0.72 | 2.74E-06 | 30 | 32 |  | 62 |
| GAREM1_219377_at | 0.73 | 1.39E-04 | 33 | 24 |  | 57 |
| LGALSL_226188_at | 0.69 | 1.08E-05 | 24 | 31 |  | 55 |
| HLF_204755_x_at | 0.71 | 7.54E-05 | 28 | 26 |  | 54 |
| SH3TC1_219256_s_at | 0.74 | 1.83E-03 | 34 | 20 |  | 54 |
| PER2_208518_s_at | 0.70 | 2.37E-04 | 26 | 23 |  | 49 |
| TMEM80_65630_at | 0.69 | 1.31E-04 | 22 | 25 |  | 47 |
| HLF_204754_at | 0.67 | 1.75E-05 | 16 | 30 |  | 46 |
| TSC22D3_208763_s_at | 0.67 | 1.86E-05 | 15 | 29 |  | 44 |
| MTERF2_225346_at | 0.69 | 2.51E-03 | 23 | 18 |  | 41 |
| BHLHE41_221530_s_at | 0.66 | 3.66E-05 | 12 | 28 |  | 40 |
| TPPP3_218876_at | 0.70 | 7.08E-03 | 25 | 10 |  | 35 |
| Mouse Data |  |  |  |  |  |  |
| Arntl 10556463 | 0.95 | 1.90E-13 | 49 | 50 | 50 | 149 |
| Npas2 10345675 | 0.93 | 8.67E-13 | 47 | 49 | 45 | 141 |
| Per3 10518781 | 0.93 | 1.25E-12 46 | 48 | 47 | 141 |  |
| Dbp 10553092 | 0.95 | 1.95E-12 50 | 47 | 43 | 140 |  |
| Per2 10356601 | 0.93 | 2.12E-11 48 | 46 | 44 | 138 |  |
| Nr1d1 10390691 | 0.92 | 1.18E-10 45 | 44 | 49 | 138 |  |
| Nr1d2 10417734 | 0.91 | 1.18E-09 | 42 | 41 | 48 | 131 |
| NA 10556487 | 0.92 | 4.16E-11 | 44 | 45 | 41 | 130 |
| Tef 10425601 | 0.86 | 2.72E-10 | 34 | 42 | 46 | 122 |
| Wee1 10556266 | 0.88 | 1.47E-09 | 38 | 39 | 37 | 114 |
| Ciart 10373452 | 0.90 | 2.59E-10 | 9 | 43 | 29 | 111 |
| Clock 10530733 | 0.82 | 3.37E-09 | 28 | 37 | 40 | 105 |
| Usp2 10584634 | 0.84 | 1.05E-08 | 33 | 35 | 33 | 101 |
| Tspan4 10558961 | 0.80 | 2.51E-09 | 24 | 38 | 36 | 98 |
| Ciart 10500272 | 0.87 | 1.39E-09 | 37 | 40 | 20 | 97 |
| Fmo2 10359582 | 0.79 | 1.44E-08 | 21 | 33 | 42 | 96 |
| Dtx4 10466304 | 0.78 | 6.36E-09 | 20 | 36 | 38 | 94 |
| Cry1 10371400 | 0.83 | 2.46E-08 | 30 | 32 | 32 | 94 |
| Lonrf3 10599192 | 0.81 | 1.79E-07 | 26 | 28 | 34 | 88 |
| Nfil3 10409278 | 0.84 | 4.90E-08 | 32 | 31 | 23 | 86 |
| Rorc 10494023 | 0.74 | 1.09E-08 | 10 | 34 | 39 | 83 |
| Tns2 10427095 | 0.83 | 1.66E-07 | 31 | 29 | 21 | 81 |
| Ypel2 10389581 | 0.82 | 1.95E-07 | 27 | 27 | 27 | 81 |
| Hlf 10389786 | 0.72 | 3.17E-07 | 3 | 25 | 35 | 63 |
| Leo1 10587211 | 0.71 | 1.38E-07 |  | 30 | 31 | 61 |
| Fbn1 10487040 | 0.74 | 1.42E-06 | 13 | 19 | 28 | 60 |
| Per1 10377439 | 0.91 | 3.00E-06 | 43 | 15 |  | 58 |
| Dapk110405693 | 0.71 | 2.82E-07 |  | 26 | 30 | 56 |
| Bhlhe41 10549276 | 0.75 | 4.60E-07 | 14 | 24 | 11 | 49 |
| Por 10526363 | 0.71 | 5.67E-07 |  | 22 | 26 | 48 |

### S8 Extra data on timing, distribution of Θ values and variable gene phase

See section S5.1 for a description of the datasets.

**Figure S3:**
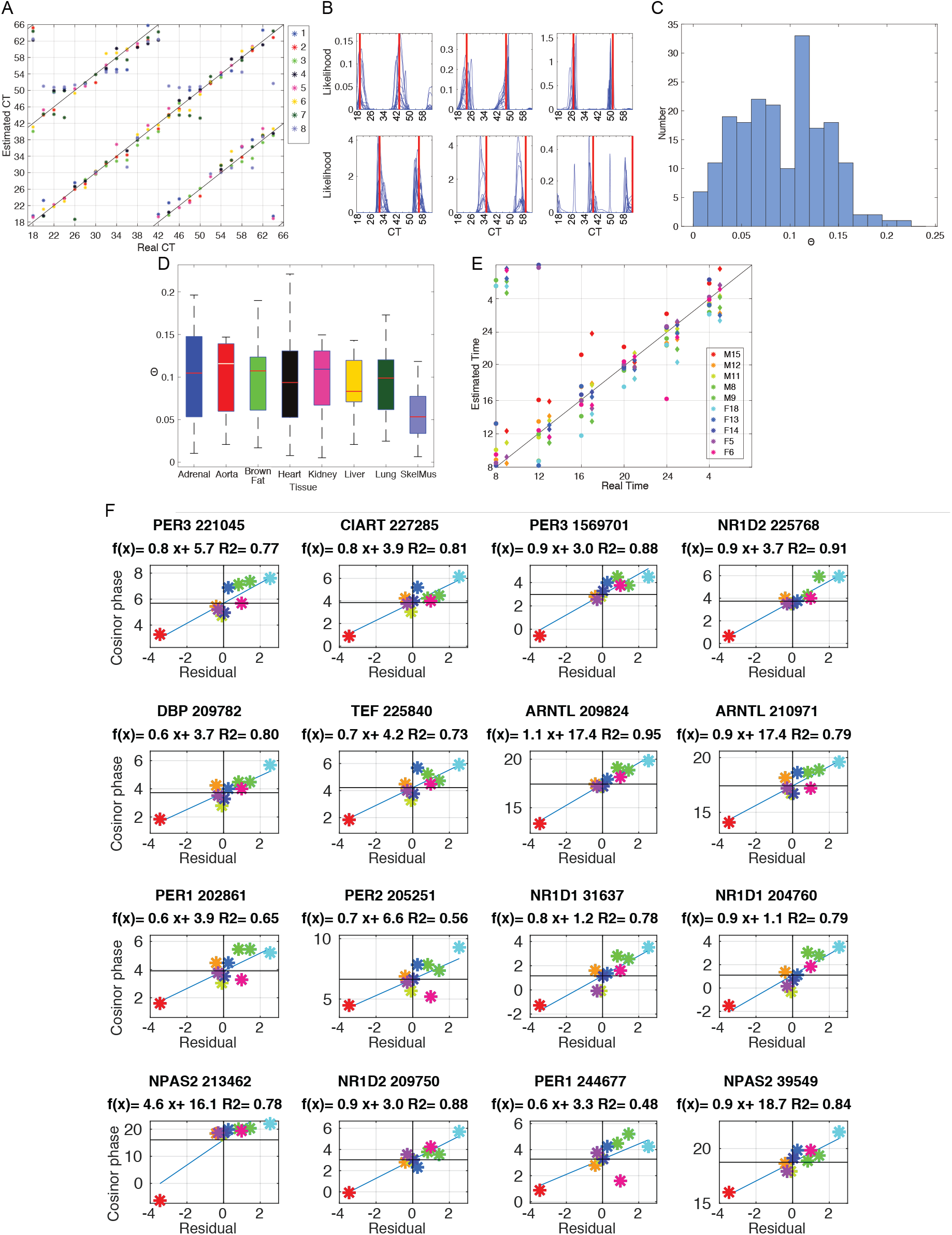
**A.** Scatter plot for actual versus predicted time for the Zhang *et al.* mouse training data (Note S5.1) using 11 probes and 8 organs. In this and the other subfigures a leave-one-tissue-out approach was used with *l*_thresh_ = −5. **B.** The likelihood functions obtained for the samples from a representative time for the mouse training data. **C.** Histogram of Θ values for the Zhang et al. data. **D.** Box plots of the distribution of Θ values for the Zhang *et al.* data for 8 of the different tissues. The Kruskal-Wallis test gives the following results for these groups: a p-value of 0.0118 for the null hypothesis that the data for each of these organs comes from the same distribution. The alternative hypothesis is that not all samples come from the same distribution. When the 8th group (Skeletal Muscle) is removed, for the other 7 groups, it gives a p-value of 0.9658. This suggests that while Skeletal Muscle has significantly lower Θs, there is no evidence that the distributions of the other seven tissues differ from each other. **E. Comparison of timings for the human training data from Bjarnason *et al.*** See Note S10.4. The filled circles show the TimeTeller estimated timing and the diamonds that for ZeitZeiger. The latter have been moved to the right by one hour of their true position to enable comparison. The data for each individual is coloured as shown in the legend. **F. Much of the estimation “error” is due to chronotype variation.** In each plot the *x*-coordinate is the difference between the time estimated by TimeTeller and the real time and the *y*-coordinate is the phase *p*_*g*_ of the indicated gene *g*. We plot the mean TimeTeller error for each individual (averaging over times) against the phase of the given gene for each individual. The horizontal line marks the mean phase of that gene. We then for each gene use the MATLAB function fit to fit a straight line *y* = *q_g_x* + *p*_*g*_ to the points as plotted. Thus, if all the “error” was due to chronotype variation the points would lie on a fitted line *y* = *q_g_x* + *p*_*g*_ which is the indicated sloping line. The closeness of the points to this line indicates that much of the “error” is due to gene-phase variation rather than TimeTeller error. Indeed, when we correct for the phase of the genes the mean absolute error for the genes (in the same order as in the figure) is 0.6423, 0.3777, 0.4075, 0.5643, 0.6928, 0.5342, 0.2705, 0.5871, 0.7500, 0.7658, 0.5657, 0.5769, 0.3858, 0.5856, 0.7813, and 0.5318 giving an average of 0.5637h. We have coloured the points to indicate which individual they correspond to (as in Figs. 1 & S3) and this demonstrates the high coherence of the (*x, y*) position for each individual across genes. There is a strong positive correlation between estimated time difference from real clock time in most of the 16 probes (and some interesting differences for some). This shows that TimeTeller can measure chronotype and predict the phase of genes such as Bmal1 (ARNTL) or REV-ERBα (NR1D1) with good accuracy if the real clock time of the sample is known. In fact, these two genes show very high correlations.

### S9 Further supplementary information about human & mouse datasets

**Figure S4:**
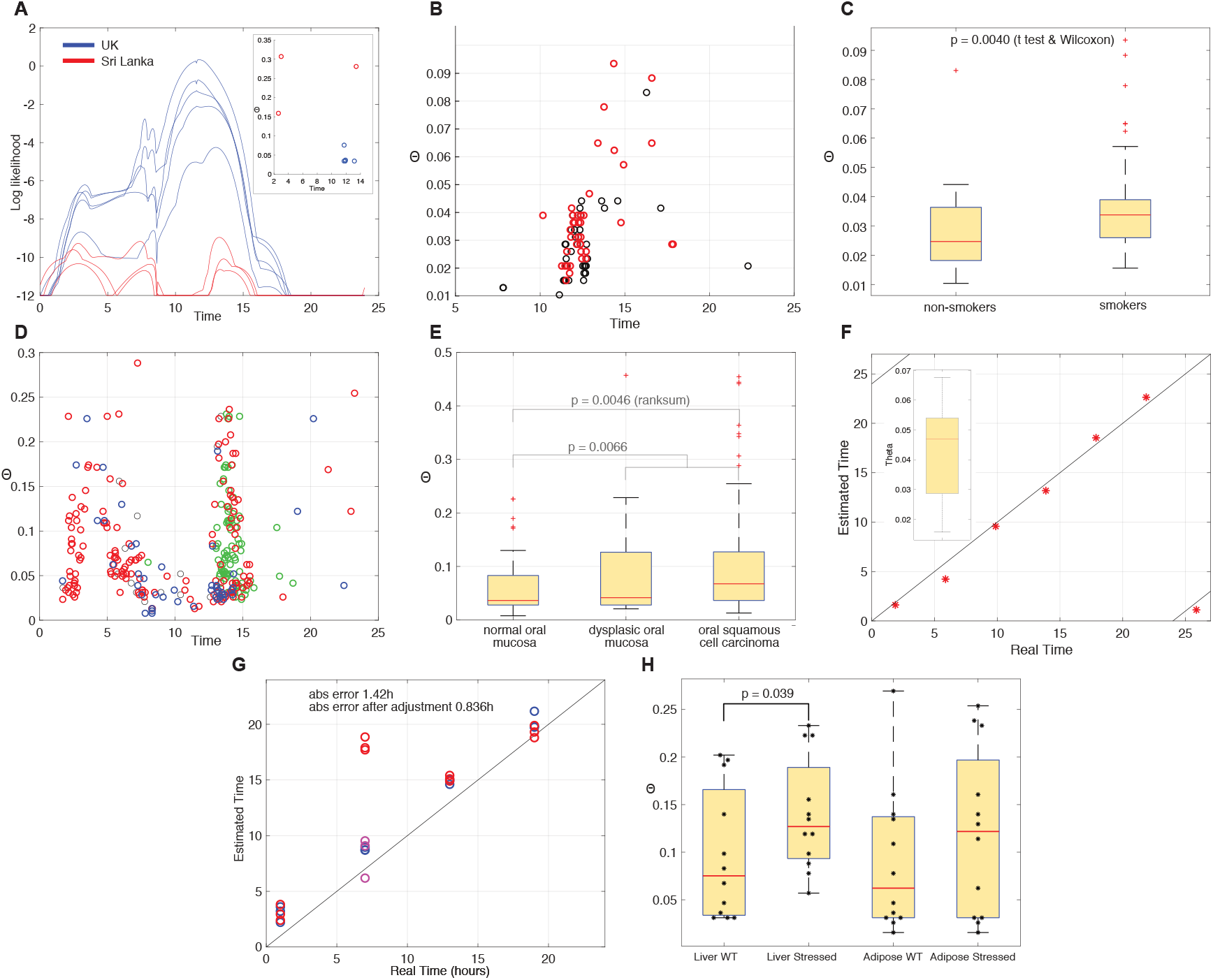
**A. Calculated log likelihood curves for the Saeed *et al.* data (*8*)** (see Note S5.3). The log likelihood curves for the 3 Sri Lankan samples (red) are very low in comparison to those for the UK data (blue). The Θ values and estimated timing are shown in the inset. The timings for the UK data (blue) appear to be realistic values reflecting our expectations. For the 3 Sri Lankan samples the Θ values are significantly larger (with two MLEs at around 2am), indicating that there is something wrong with these samples. The reasons behind this can only be speculated upon. However, this suggests that TimeTeller may have a useful function in terms of quality control. The log threshold *l*_thresh_ used here is −12. **B,C. TimeTeller’s analysis of the Boyle *et al.* data (*9*)**. **B.** A scatter plot of Θ values against estimated time (non-smokers (black) and smokers (red)). Most of the estimated times are between 8am and 4pm. The majority of the 79 non-smoker and smoker samples have Θ < Θ_GCF_ = 0.1, which are in the GCF range. C. However, the boxplot shows that the median Θ for non-smoker samples is smaller than the median Θ for the smoker samples, with a significant Wilcoxon test statistic of *p* = 0.004. These results provide further evidence that human datasets show reliable well-defined likelihood functions and that one can observe differences in Θ between WT and perturbed clocks. The log threshold *l*_thresh_ used here is −12. **D,E. Timing and Θ values for the Feng *et al.* dataset (*10*). D. Estimates for the timing MLE *T* plotted against Θ values from TimeTeller for the Feng *et al.* dataset (*10*)**. This uses *l*_thresh_ = −12. The estimates for the normal data (blue) are generally between 9 am and 6 pm, where the majority of the estimates outside of this time range are for cancer samples (red). The green points indicate the times of the second peak for those samples with *T* < 7. This suggests that the mistimed samples are primarily so because the wrong peak has a higher likelihood. **E. The distribution of Θ values by group.** There is a highly significant difference in median values between the cancer group (*n* = 167) and both the normal group (*n* = 45) and the combined normal/dysplasia group (*n* = 62), with the p-values indicated. **F. Scatter plot of real times against estimated times for the Martelot *et al.* dataset (*5*)**. (Inset) Boxplot of the corresponding Θ values. **G,H Timing and Θ distribution for the Barclay *et al.* data on sleep stress.** See Sects. S5.2 & S9.1.

#### S9.1 Timing for the Barclay *et al.* dataset

Barclay *et al.* (*7*) published a dataset that uses all WT mice, but half of the mice were sleep deprived when they would usually be sleeping. See Note S5.3. This is an interesting example where the clock is perturbed rather than structurally dysfunctional. In this dataset all the mice are WT, but half of them were sleep deprived when they would normally be sleeping. See Note S5.2 for more details.

Time-Teller accurately estimated the time of the normal condition liver samples, shown as the blue circles on the left plot in Fig. S4(G). The Θ values for these samples have a median Θ value around 0.03 (Fig. S4(H)). There is also a good level of estimation accuracy within the normal sleep schedule group for the adipose data, with the exception of an ~10 hour error for three data points for the adipose tissue at 07.00. However, this substantial error can be corrected by using the second peak of the likelihood curve for these samples.

For the sleep stressed group, some samples are inaccurately predicted, as shown on the right plot in Fig. S4(G). The Θ values for the time stressed group are generally higher, as shown in the boxplot in Fig. S4(H). As sleep stressed clocks are not technically classed as dysfunctional (*14*), we would not expect the Θ values to as large as with KO data. There is however, a notable difference of Θ values for the normal and sleep deprivation groups, which is significant in the liver data (*p* = 0.02) and less significant in the adipose date (*p* = 0.1).

This suggests that sleep stress actually changes circadian gene expression to a profile different than any gene expression profiles that occur in normal sleep-wake conditions. This provides evidence that a chronic change to the sleep-wake schedule results in abnormal circadian gene expression patterns, and not just a change in body time.

#### S9.2 Comparing rhythmic datasets

Because of the nature of the TimeTeller algorithm we were unable to find a normalisation technique to allow comparison of data from different microarray technologies with our training data without removing the key multidimensional aspects determining functionality. The time-dependent expression ratios between the genes in the rhythmic expression profile varied significantly from one technology to another. For example, we studied circadian timecourse data from two independent datasets comparing the raw expression in the liver of mice from Hughes *et al.* (*15*) (Affymetrix Mouse Genome 430 2.0 GeneChip) and Zhang *et al.* (Affymetrix MoGene 1.0ST GeneChip) after fRMA normalisation. These use two different Affymetrix genechips. Different microarrays use slightly different probe designs and quantify the expression differently because of this. The time-dependent expression ratios and probe dependent differences in mean intensity and amplitude in most probes was apparent. With the advent and dominance of RNA-seq we expect this problem to be largely solved.

### S10 Further supplementary information about the REMAGUS data set

#### S10.1 Patients’ characteristics

**Table S2:** Patients’ characteristics. While the cause of death for 4 patients is unknown, for the others it was breast cancer Abbreviations: LVI, lymphovascular invasion; pCR, pathological complete response. **n = number of patients. Missing data: HER2-CT alone: Menopausal status: 2, Nodal status: 1, Grade: 4, PR status: 1, pCR: 2. HER2-CT+celo: Grade: 1, LVI 1,pCR: 2. HER2+ CT alone: Nodal status:1,Grade: 1, pCR: 1. HER2+ CT + trastuzumab: Grade: 1, LVI : 1.

|  |  | HER2 negative |  | HER2 positive |  |
| --- | --- | --- | --- | --- | --- |
|  |  | CT alone<br>n = 77 (%) | CT + celocoxib<br>n = 69 (%) | CT alone<br>n = 39 (%) | CT + trastuzumab<br>n = 41 (%) |
| Age (years) | < 40 | 16(21) | 15(22) | 9(23) | 9(22) |
|  | 40-49 | 30(39) | 32(46) | 17(44) | 13(32) |
|  | ≥ 50 | 31 (40) | 22(32) | 13(33) | 19(46) |
| Menopausal status. | No | 48(62) | 48 (70) | 29(74) | 27(66) |
|  | Yes | 27(35) | 21(30) | 10(26) | 14(34) |
| Clinical tumour size | T2 | 41(53) | 45(65) | 18(46) | 17(41) |
|  | T3 & T4 | 36(47) | 24(35) | 21(64) | 24(58) |
| Clinical nodal status* | N0 | 32(42) | 28(41) | 13(33) | 18(44) |
|  | N1, N2 & N3 | 44(57) | 41(59) | 25(64) | 23(56) |
| Histological type | Ductal | 67(87) | 56(81) | 37(95) | 37(90) |
|  | Lobular | 7(9) | 7(10) | 1(2.5) | 1(3) |
|  | Others | 3(4) | 6(9) | 1(2.5) | 3(7) |
| SBR grade* | I | 10(14) | 5(7) | 0(0) | 0(0) |
|  | II, III, IV | 63(86) | 63(92) | 38(100) | 40(99) |
| LVI | No | 66(86) | 58(84) | 30(77) | 27(66) |
|  | Yes | 11(14) | 10(14) | 9(23) | 10(24) |
| ER | Negative 74 | 25(32) | 28(41) | 17(44) | 18(44) |
|  | Positive 119 | 52(68) | 41(59) | 22(56) | 23(56) |
| PgR* | Negative | 40(52) | 40(58) | 24(62) | 24(59) |
|  | Positive | 36(47) | 27(39) | 15(38) | 17(41) |
| Triple negative | Yes | 1(1) | 2(3) | 13(33) | 16(39) |
|  | No | 74(99) | 67(97) | 26(67) | 25(61) |
| HR (4 classes) | ER-/PR- | 24(31) | 28(41) | 15(38) | 17(41) |
|  | ER+/PR- | 17(22) | 12(17) | 9(23) | 7(17) |
|  | ER-/PR+ | 35(45) | 27(39) | 2(5) | 1(2) |
|  | ER+/PR+ | 1(1) | 2(3) | 13(33) | 16(39) |
| Neoadjuvant Treatment | No celocoxib | 77(100) | 12(17) | 39 | 41(100) |
|  | Celocoxib | 0(0) | 57(83) | 0 | 0 |
|  | No trastuzumab | 77(100) | 69(100) | 37(95) | 2(5) |
|  | Trastuzumab | 0(0) | 0(0) | 2(5) | 39(95) |
| pCR | No | 67(87) | 56(81) | 32(82) | 31(76) |
|  | Yes | 8(10) | 11(16) | 6(15) | 10(24) |
| Death | No | 61(81) | 50(72) | 31(82) | 35(85) |
|  | Yes | 14(19) | 19(28) | 7(18) | 6(15) |

#### S10.2 Survival analysis for disease free survival (DFS) & overall survival (OS)

Recall that the meaning of the hazard rate HR for the multivariate analysis is that if we hold the other prognostic factors constant at their mean HR values, the ratio of the hazard of an individual with Θ^’^ to that of one with Θ is HR(Θ’-Θ). Thus, if HR < 1 then the hazard decreases with increasing Θ. Also recall that we scale Θ by 10 and use the modified hazard rate HR^1/10^ which is the hazard rate for 10 × Θ or equivalently the ratio for a Θ increase of 0.1. This makes HR^1/10^ sensibly comparable with the hazard ratio values of the other prognostic strata. The HR values for Θ without this scaling are very small. The p-value for the alternative hypothesis HR < 1 is the same as for HR^1/10^ < 1.

**Table S3:** Results from univariate & multivariate analysis of disease-free survival and overall survival using the Cox Proportional Hazards Model. For each prognostic stratum the HR^1/10^, its confidence limits, the number of events and individuals, and the corresponding p-value are shown. The p-values associated with each factor concern the HR for that factor where one compares the hazard for Θ against the hazard for the alternative, conditional on the other prognostic factors being the same. The prognostic groups shown for multivariate analysis are those for which this univariate analysis p-values is ≤0.05. This analysis was implemented using the R package coxph. The other factors included in the multivariate analysis were ER, PR and HER2 status, tumour grade and size of tumour.

|  | DFS |  |  |  | OS |  |  |  |
| --- | --- | --- | --- | --- | --- | --- | --- | --- |
| Univariate analysis |  |  |  |  |  |  |  |  |
| Stratum | HR<sup>1/10</sup> | Conf. Lims. | n=/events | p-value | HR<sup>1/10</sup> | Conf. Lims. | n=/events | p-value |
| All | 0.46 | 0.27–0.80 | 205/58 | 0.005** | 0.4683 | 0.24-0.90 | 209/41 | 0.01** |
| ER+ | 0.57 | 0.31–1.02 | 127/37 | 0.06 | 0.40 | 0.14-1.16 | 129/18 | 0.05* |
| ER- | 0.37 | 0.14–0.98 | 78/23 | 0.05* | 0.46 | 0.19-1.11 | 80/23 | 0.06 |
| PR+ | 0.58 | 0.28–1.18 | 90/22 | 0.1 | 0.60 | 0.20-1.79 | 91/10 | 0.3 |
| PR- | 0.46 | 0.22–0.96 | 112/36 | 0.04* | 0.41 | 0.18-0.94 | 115/31 | 0.02* |
| HER2+ | 0.24 | 0.05–1.13 | 74/14 | 0.07 | 0.21 | 0.03-1.35 | 75/11 | 0.03* |
| HER2- | 0.55 | 0.32–0.95 | 131/46 | 0.03* | 0.54 | 0.27-1.07 | 134/30 | 0.05* |
| pCR+ | 0.12 | 0.004–3.38 | 32/4 | 0.2 | 0.11 | 0.004-2.96 | 32/4 | 0.09 |
| pCR- | 0.57 | 0.34–0.95 | 173/56 | 0.03* | 0.49 | 0.24-0.99 | 174/35 | 0.03* |
| grade=1/2 | 0.71 | 0.34–1.48 | 90/24 | 0.4 | 0.69 | 0.26-1.85 | 90/14 | 0.4 |
| grade = 3 | 0.34 | 0.15–0.75 | 109/34 | 0.008* | 0.28 | 0.10-0.77 | 112/25 | 0.002* |
| size = 2 | 0.50 | 0.25–1.03 | 110/27 | 0.06 | 0.27 | 0.06-1.2 | 110/11 | 0.04* |
| size = 3/4 | 0.51 | 0.26–1.0 | 108/33 | 0.052 | 0.40 | 0.14-1.12 | 108/19 | 0.05* |
| nodal stat. 1 | 0.35 | 0.17–0.75 | 125/41 | 0.006* | 0.35 | 0.14-0.87 | 127/28 | 0.007** |
| no LVI | 0.48 | 0.26–0.90 | 162/43 | 0.02* | 0.52 | 0.25-1.09 | 165/29 | 0.06 |
| LVI | 0.42 | 2.38–0.14 | 38/15 | 0.1 | 0.41 | 0.11-1.52 | 39/11 | 0.1 |
| TN | 0.57 | 0.2025–1.6 | 44/17 | 0.3 | 0.65 | 0.26-1.66 | 46/18 | 0.3 |
| not TN | 0.48 | 0.26–0.87 | 161/43 | 0.02* | 0.34 | 0.13-0.91 | 163/23 | 0.01** |
| Multivariate analysis |  |  |  |  |  |  |  |  |
| All | 0.46 | 0.27-0.80 | 205/58 | 0.005** | 0.42 | 0.21–0.86 | 209/39 | 0.18* |
| ER- | 0.54 | 0.29-0.98 | 127/37 | 0.043 |  |  |  |  |
| PR- | 0.41 | 0.18-0.94 | 112/34 | 0.035* |  |  |  |  |
| HER2- | 0.53 | 0.30-0.95 | 131/44 | 0.033* |  |  |  |  |
| HER2+ | 0.22 | 0.049-0.99 | 74/14 | 0.048* | 0.19 | 0.035–1.1 | 75/11 | 0.58 |
| pCR- | 0.53 | 0.31-0.91 | 173/54 | 0.22* | 0.48 | 0.22–1.2 | 173/33 | 0.071 |
| Grade 3 | 0.36 | 0.16-0.82 | 109/34 | 0.15* | 0.28 | 0.096–0.82 | 112/25 | 0.02* |
| Nodal Stat. 1 | 0.30 | 0.13-0.69 | 124/39 | 0.005** | 0.21 | 0.06–0.7 | 127/26 | 0.011* |
| no LVI | 0.48 | 0.26-0.89 | 162/43 | 0.021* |  |  |  |  |
| not TN |  |  |  |  | 0.35 | 0.13–0.95 | 163/23 | 0.04* |

#### S10.3 Real *vs*. estimated times for REMAGUS breast cancer samples, comparison with ZeitZeiger, correlation with PCNA, & the distribution of Θ values for the main prognostic strata

**Figure S5:**
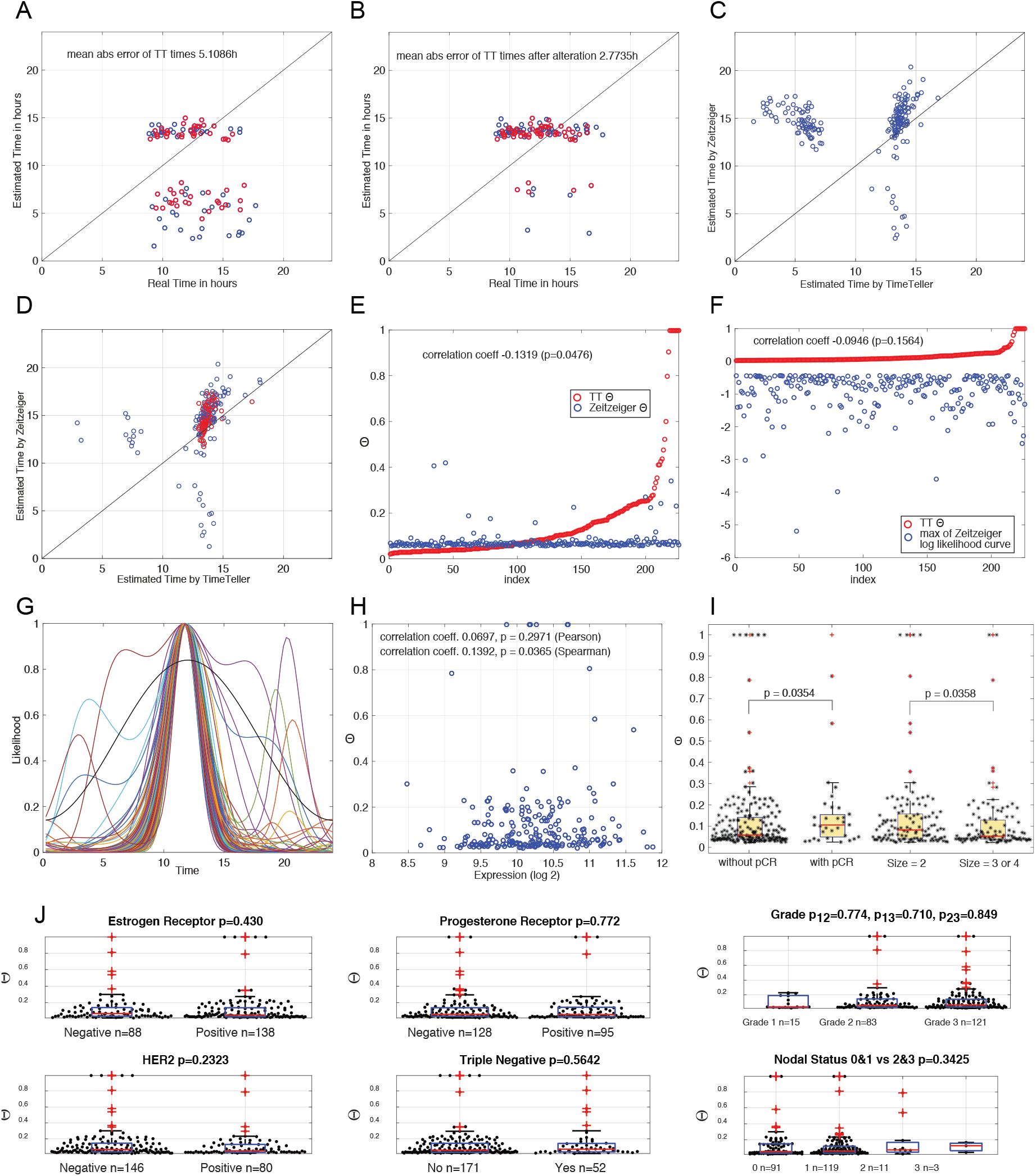
(A) Scatter plot showing real *vs*. estimated times for those 108 REMAGUS breast cancer samples that had timing information. This uses *l*_thresh_ = −12. All of the tumour biopsies were performed between 8:45 and 17:45. The markers are coloured by Θ ≤ Θ_GCF_ (red) and Θ > Θ_GCF_ (blue) where Θ_GCF_ = 0.1. There is a clear “daytime cluster” around midday on both axes where most of the estimates lie. Time estimates for samples with a functioning clock have a mean error of < 3 hours. **(B) Second peak correction.** This plot shows the timing scatter plot if for those sample outside the “daytime cluster” with a clear second peak one uses the second peak to define the estimated time. The proportion of badly timed samples decreases to 9%. **(C) Scatter plot of the estimated times for the REMAGUS data comparing TimeTeller and ZeitZeiger.** Both show a dominant midday cluster and a number of samples with unreasonably early timing, TimeTeller having more than ZeitZeiger. **(D) As (B) but with second peak correction.** The mistimed points appear corrected if the second peak is used to define time *T*. The red points show timing for the samples where TimeTeller gave seemingly unreasonable times and we replace the estimated time with the time given by the second peak. With this adjustment both algorithms get realistic times for more than 90% of the data that has a non-flat likelihood curve. For the 108 samples with timing they get comparable mean absolute errors: 2.9280h for ZeitZeiger and 2.7735 for TimeTeller. On the other hand, ZeitZeiger appears to get a better spread of estimates over the working day (i.e. between 8.45 and 17.45 during the “daytime cluster” shown in A. - E.) and less samples with unreasonable times. The Spearman correlation between the timings in the main cluster after second peak correction is 0.6461 (*p* ≈ 0). **(E) A comparison of TimeTeller Θs from all REMAGUS samples with those from ZeitZeiger.** See Note S10.4. For ZeitZeiger we take the ZeitZeiger likelihood curve and then define Θ as for TimeTeller. The samples are ordered in increasing TimeTeller Θ and the TimeTeller Θ and the ZeitZeiger Θ for a given sample are plotted with the same *x*-coordinate. We see that although there is a very small negative correlation, there is essentially no relationship between the two and the variation in the ZeitZeiger across all samples is very small compared to that of TimeTeller. **(F) Using the likelihood maximum for ZeitZeiger.** As in (E) except that the ZeitZeiger Θs are replaced by the maximum value of the ZeitZeiger loglikelihood function as described in Note S13.2. **(G) A plot of all the ZeitZeiger centred likelihood ratio functions.** These show much less variation than the TimeTeller ones (Fig. S2)which explains the relative uniformity of the ZeitZeiger Θ across samples in (E). **(H) Scatter plots of Θ against the expression of the genes PCNA and CKS2 for the REMAGUS data.** The p-values are for testing the alternative hypothesis of a non-zero correlation against the null hypothesis of no correlation. **(I,J) Θ distributions against prognostic strata.** (I) Boxplots showing distributions of values for the prognostic strata corresponding to tumour size and pathologic Complete Response (pCR) after the administration of neo-adjuvant chemotherapy. (J) Boxplots showing distributions of values for the other main prognostic strata: Estrogen receptors, Progesterone receptors, HER2 receptors, Triple Negative status, Grade, and Nodal Status.

#### S10.4 Comparison of dysfunction metric with ZeitZeiger for the REMAGUS data

It was suggested to us that we should compare our results on the REMAGUS data with Zeitzeiger ((*16*), Note S13.2) using Zeitzeiger’s likelihood function, noting that Zeitzeiger’s authors never claimed it could calculate dysfunction in individual samples. However, we have now included an analysis in Fig. S5(E,F). The only way we could see to do this was by using the Zeitzeiger likelihood in a similar way to TimeTeller’s.

The Bjarnason *et al.* data was used to train ZeitZeiger which seemingly accurately estimated the times of the training data in a cross-validation study as seen in Fig. S3E (parameters sumsabsv = 1 and nSpc = 2 were chosen). We used the ZeitZeiger package (https://github.com/hugheylab/zeitzeiger). The ZeitZeiger timings and likelihoods were then calculated for the rhythmic expression profiles of the REMAGUS data.

As well as comparing Θ values (Fig. S5E) we compared TimeTeller Θ values with the maximum value of the ZeitZeiger likelihood curve (Fig. S5F). This is because in (*16*) it was suggested that this might be characteristic of a dysfunctional clock as they observed that in four of seven datasets studied in that paper, the likelihood of the predicted timing was significantly lower in mutant than in wild-type.

#### S10.5 Relevance of Venet *et al.*

The paper (*17*) considers the common technique of clustering patients using gene expression from a candidate prognostic gene set asking for specific mis-expression of the genes in this set. The key point in the paper is that for a substantial proportion of 47 proposed multi-gene breast cancer markers which apparently showed a significant link to survival, this link was markedly weakened if the expression of the genes in a list called meta-PCNA were removed. These meta-PCNA genes are consistently expressed when the gene PCNA is expressed in normal tissues and consistently repressed when PCNA is repressed. They pointed out that this is also behind the tendency of random markers to show an association because PCNA expression is associated with an increased cell proliferation.

This raises the question of whether such an observation applies to our dysfunction metric Θ and its strong association with survival in the REMAGUS data. There are three reasons why this does not apply.

Firstly, none of the genes we use are in the meta-PCNA list. Secondly, these markers are all defined by whether their genes are under- or over-expressed and therefore the correlating effect of the meta-PCNA genes has to do with whether they are under- or over-expressed. Our marker is completely different in nature as it looks to see if the clock genes are in the correct relationship with the other clock genes being considered and so it cannot be correlated with meta-PCNA in the way that the studied markers are. Thirdly, we have analysed the behaviour of the two dominant genes in the meta-PCNA list (i.e. PCNA and CKS2) and showed that in the REMAGUS data there is no significant correlation between Θ values and expression of these genes (see Fig. S5H). For PCNA, ρ = 0.07 (Pearson, *p* = 0.3) & 0.137 (Spearman, *p* = 0.037), while for CKS2 *ρ* = 0.03 (Pearson, *p* = 0.6) & 0.03 (Spearman, *p* = 0.64)

#### S10.6 Which aspects of TimeTeller likelihood curves contain biological information?

The metric Θ is constructed so as to assess both the confidence interval of the MLE primary peak timing *T* and the competition from alternative peaks (Note S2). A good way to get an understanding of these matters is to consider the centred likelihood ration curves in Figs. S4 and S5. These have been centred in that the primary peak has been moved to *t* = 12 so that the curves can be easily compared. The Θ associated to each curve is the proportion of time it spends above the curve *C*(*t*|*T*) defined in Note S2. These show a clear difference in structure between the samples with GCF and those with larger Θs. This raises the question of which aspects of the likelihood curves can identify interesting aspects of disease and, in particular, to what extent the presence/absence of a second peak is what is determining the significant hazard ratios found in the REMAGUS data. Consequently, we repeated the 0 < Θ ≤ 0.1, and with a single peak (102 out of 109). We found that for this population Θ has a mean *HR*^1/10^ of 0.13 (*p* = 0.11). Moreover, if we define a threshold at Θ_*T*_ = 0.06 and only consider these samples, the Kaplan-Meier survival plot comparing Θ < Θ_*T*_ with Θ ≥ Θ_*T*_ shows a difference in 10 year survival of nearly 20% (*p* = 0.06). Since there is only one peak for these samples, Θ is a direct estimate of the confidence interval as in the classical statistical context (Note S4). This shows that there is interesting biological information in the size of this confidence interval. Then we took all the samples with two peaks and analysed this similarly. Again, we found a mean *HR*^1/10^ below 0.25 suggesting that we can extract interesting information from this population, though this estimate is not statistically significant. We conclude That for data with a single peak the classical construct provides interesting biological information and the extended one used here that deals with second peaks adds to this.

### S11 Supplementary information about synthetic datasets: 1. Local PCs versus global PCs, 2. Θ dependence on the efficiency of a Bmal1 knockdown

**Figure S6:**
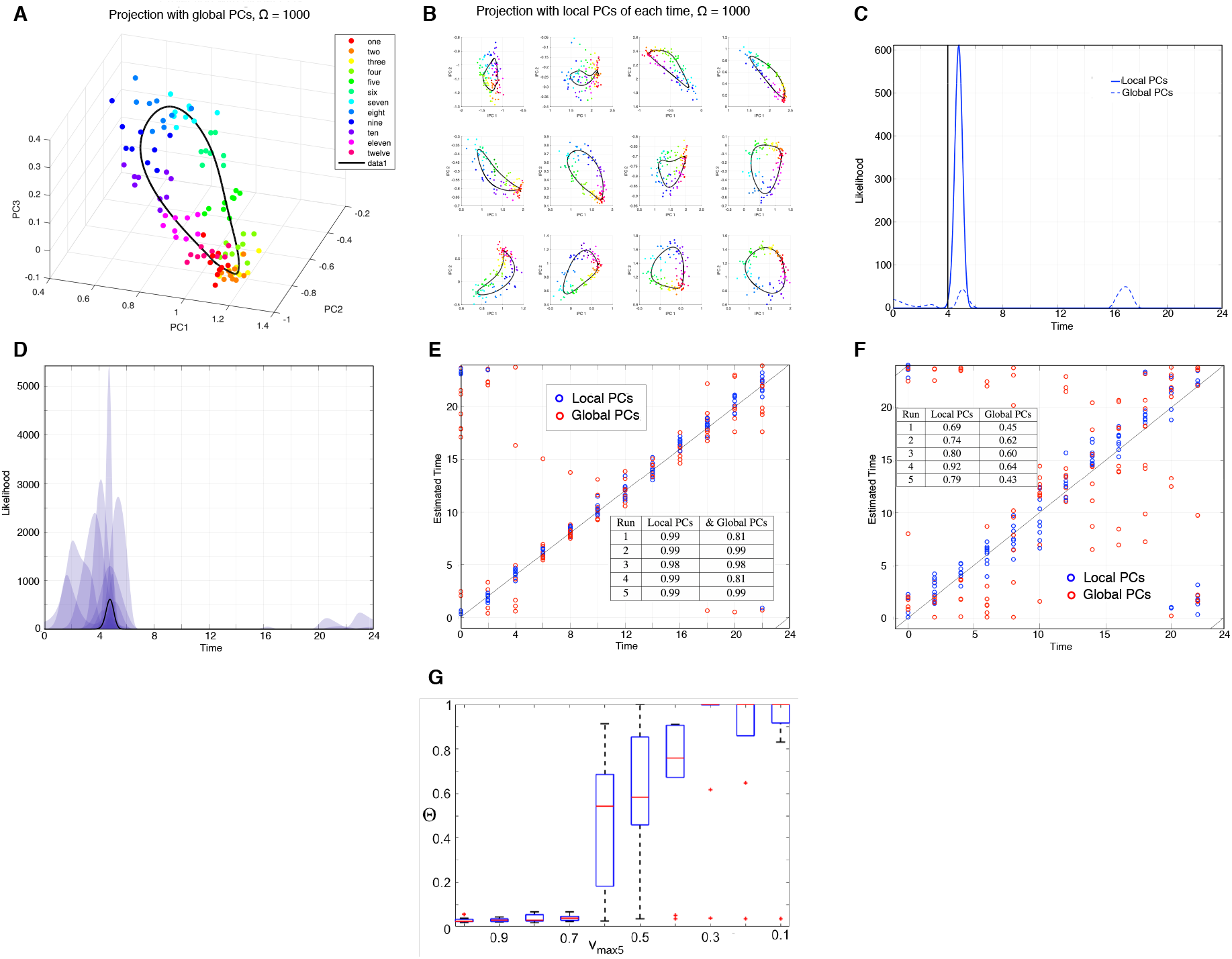
**A-F. The construction of the TimeTeller probability model and likelihood function uses projections calculated locally (local PCs) which are then combined.** We compared the effectiveness of this local approach to the simpler one using a single principal component projection of all the training data (global PCs). To do this we used simulated data from the stochastic Relogio model (Note S12.1) using systems sizes Ω = 1000 and Ω = 100 and 12 time points around the day. The latter system size produces particularly noisy data. (A) A global PC projection of the data when Ω = 1000. (B) The 12 local PC projections. (C) Example likelihood curves for the same data point for global and local PCs. The global PC likelihood is shown with the dashed line and the local PC likelihood is shown in bold, for the same training and test data. The black vertical line shows the true time *T* of the sample. Notice the improvement in accuracy when using local PCs and the MLE peak at *T* + 12 for the global PCs. (D) Plot showing how individual local PC likelihoods are averaged to produce the combined likelihood. The likelihood curves for the local PCs are (geometrically) averaged, to get the overall likelihood (Note S1.2). This helps to overcome problems with symmetry as described in Fig. S1 and seen in (C). (E) Scatter plot showing estimated versus real time for low noise (Ω = 1000) simulations, using local and global PCA. Correlation is generally very high, with some obvious symmetry issues with the global PC estimations. The red point at (8, 13) is the estimate resulting from the dotted likelihood in (C). The inset table shows *R*^2^ values for linear fit to *x* = *y* of estimated versus real time, for Ω = 1000 simulated data. Values are generally accurate due to low noise. *R*^2^s are similar except for runs 1 and 4, where the accuracy of the local PC method is higher than for global PC method. (F) As (E) but for the high noise Ω = 100 simulations. The training and test data are identical. From (E) and (F) and the inset tables we observe that the method using local PCs is more accurate than the method using global PCs. The inset table is as for (E) but for Ω = 100. Values vary due to high noise in the simulated data. *R*^2^s for local PCs are significantly higher than for global PC. **(G) Θ dependence on the efficiency of a Bmal1 knockdown.** We investigated how Θ increases as we increase the efficiency of a Bmal1 knockdown using synthetic data from the stochastic Relogio model with intermediate systems size Ω = 500, giving a moderate level of stochasticity. Bmal1 is crucial for the proper functioning of the circadian clock (*18*). The knockdown of Bmal1 is modelled by decreasing the rate *V*_*max*_ 5 of Bmal1 transcription from the WT value of 1 to the full knock-down value of 0.1, in steps of 0.1. For each of ten stochastic simulations, 10 equally spaced time-points were extracted as the test sets. The box plots show that there is an (almost) monotonic relationship between Θ and the change in the *V*_*max*_ 5 parameter and that, as *V*_*max*_ 5 falls below 0.7, the functioning of the clock is severely affected.

### S12 Stochastic modelling

There have been a number of stochastic models of the circadian clocks (e.g. (*19–21*)). In particular, an early detailed stochastic mammalian model of circadian rhythms was published by Forger *et al.* (*22*), who adapted his previous 74 equation ODE model. Instead of attempting to replicate this existing stochastic model of mammalian circadian rhythms, we have created a stochastic version of the Relogio model (*23*) with the adaptation based on methods in (*19*). This stochastic model incorporates a system size parameter, Ω. We consider it has units L/nM and a value in the range 100-1000 based on typical cell size estimates (*24*). The individual trajectories were calculated in MATLAB, using an implementation of the Gillespie algorithm or the pc-LNA (*24*).

#### S12.1 The stochastic Relogio model

The stochastic Relogio model has 44 reactions using 19 variables and 72 parameters, where parameters with units of concentration are scaled with system size parameter accordingly. As Ω → ∞, the stochastic solution converges to the ODE solutions. The 44 reaction rates for the stochastic Relogio model are shown in Note S12.1. In order to generate enough trajectories to estimate a distribution in a reasonable time the pc-LNA was used (*24*).

##### ODE

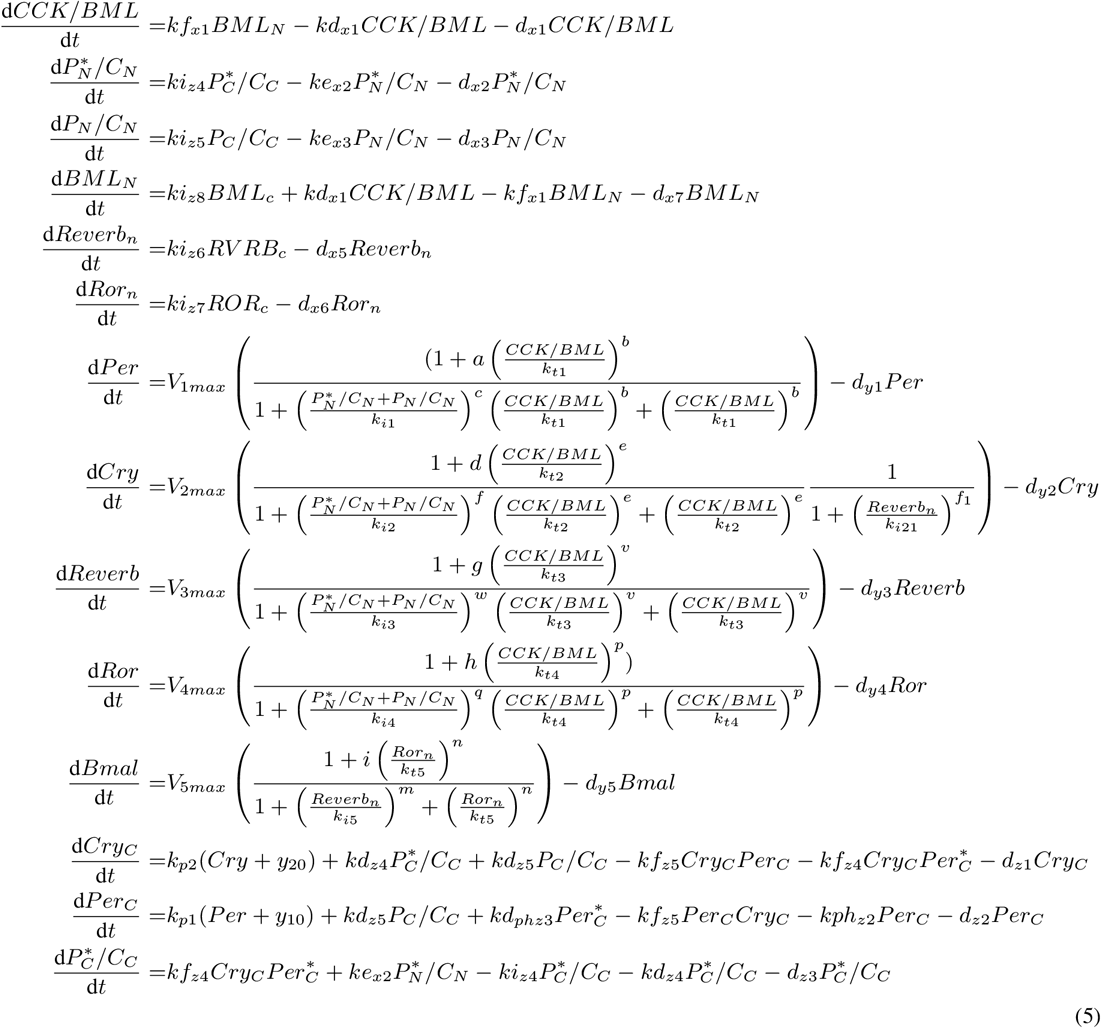

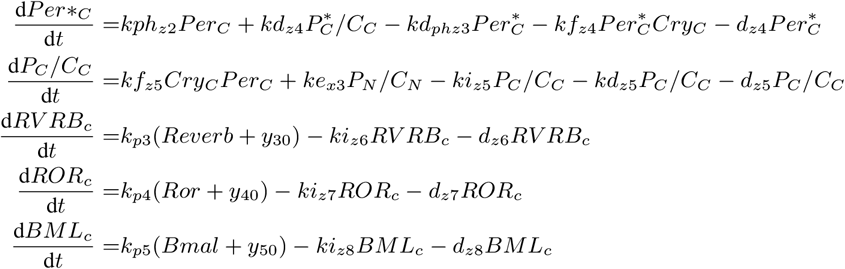

##### Stochastic model

19 state variables. 44 reactions. 72 parameters.

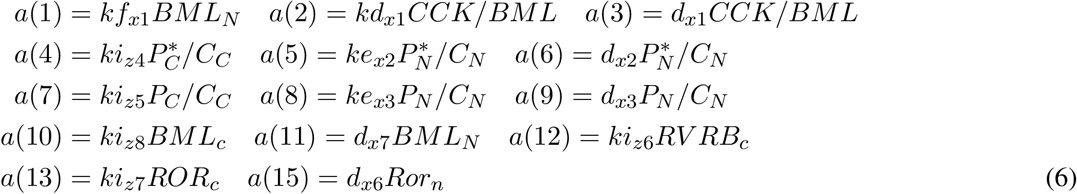

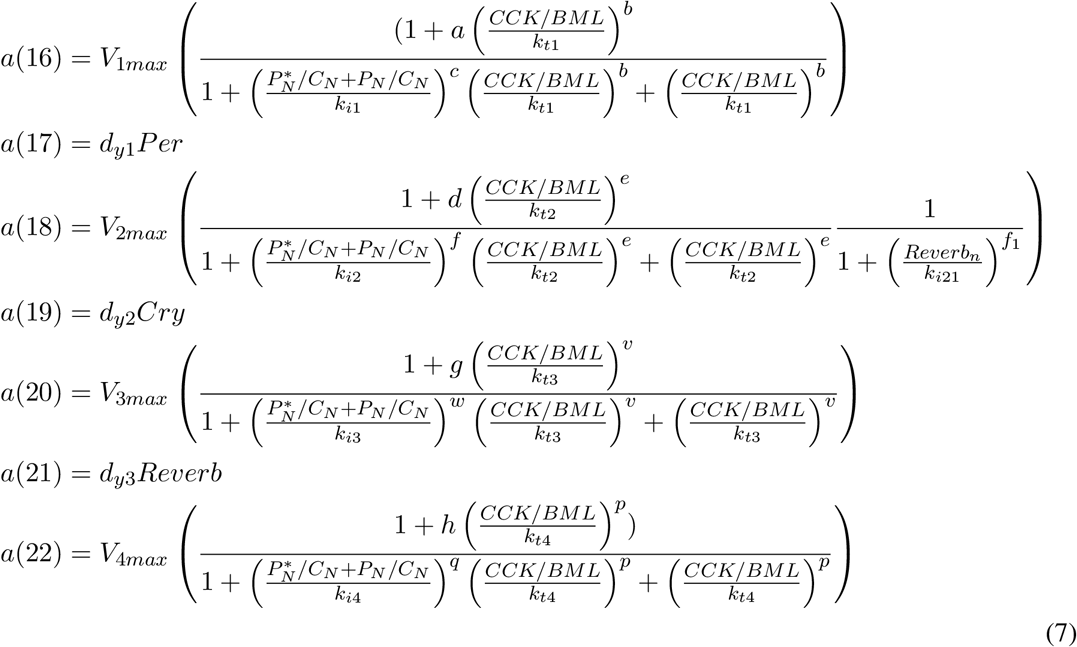

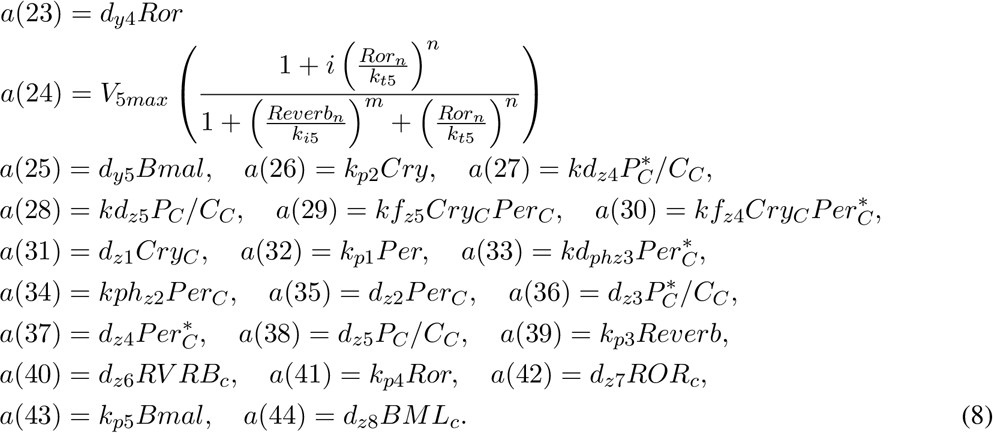

##### Parameters

**Table S4.**
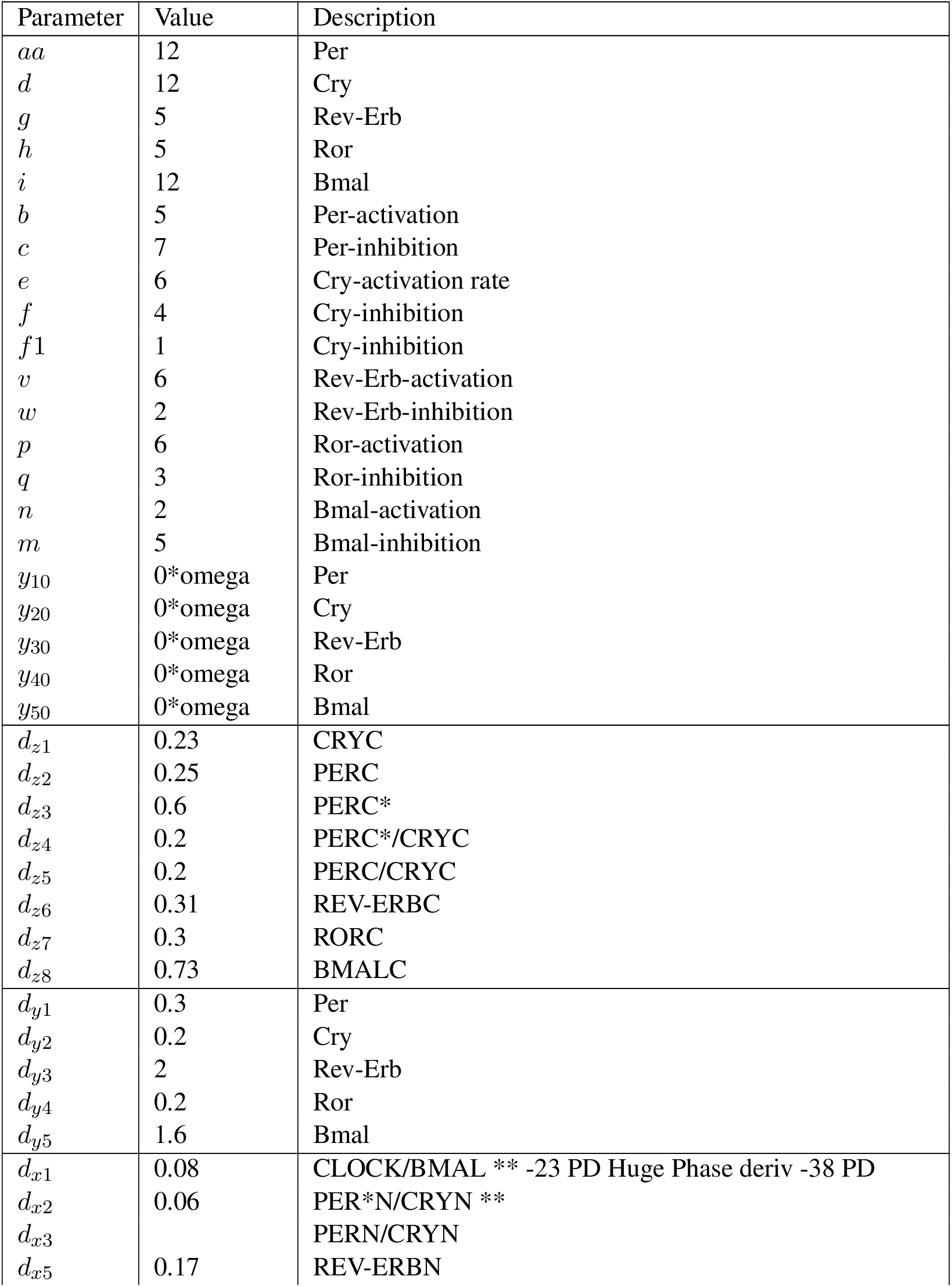
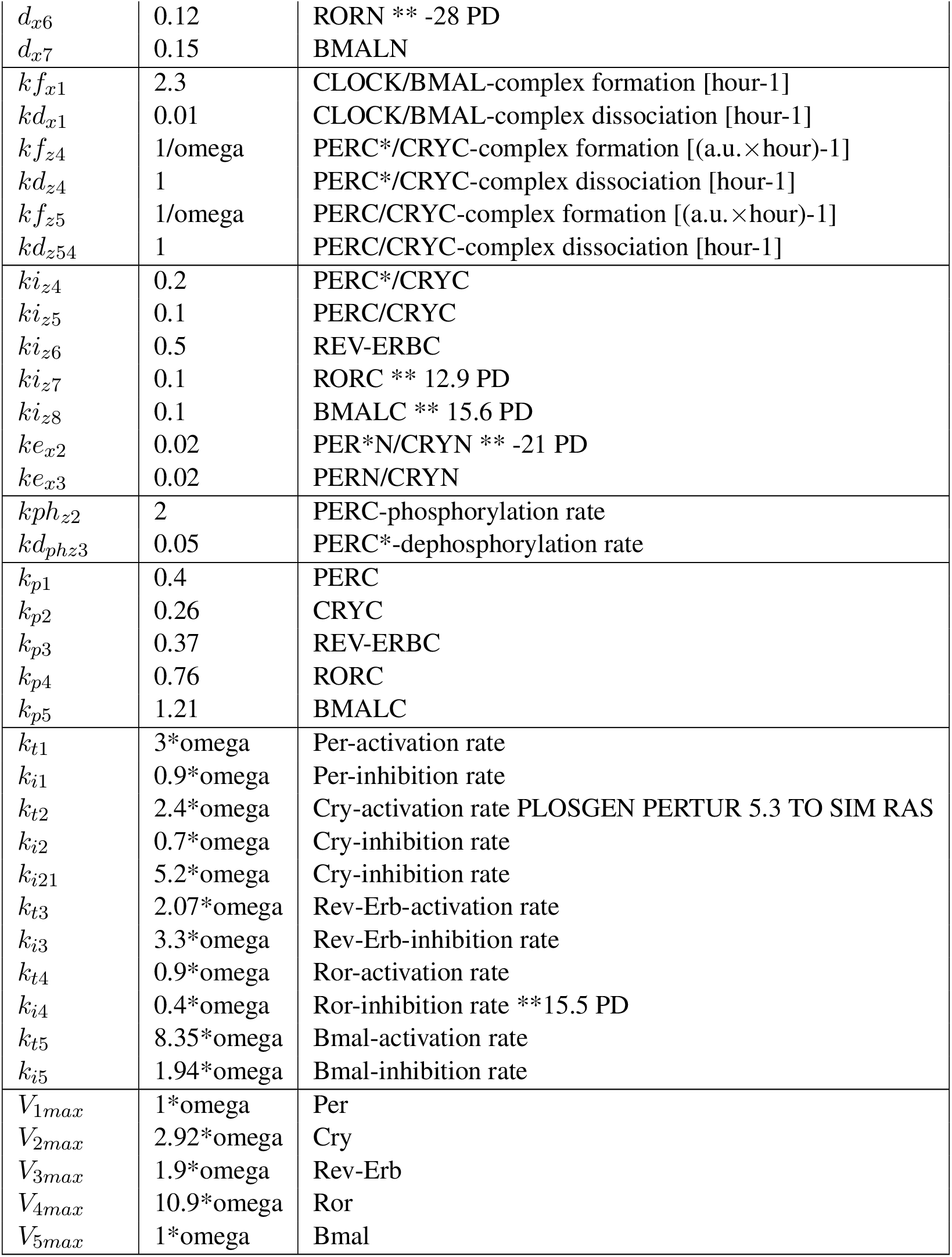

| Parameter | Value | Description |
| --- | --- | --- |
| $aa$ | 12 | Per |
| $d$ | 12 | Cry |
| $g$ | 5 | Rev-Erb |
| $h$ | 5 | Ror |
| $i$ | 12 | Bmal |
| $b$ | 5 | Per-activation |
| $c$ | 7 | Per-inhibition |
| $e$ | 6 | Cry-activation rate |
| $f$ | 4 | Cry-inhibition |
| $f1$ | 1 | Cry-inhibition |
| $v$ | 6 | Rev-Erb-activation |
| $w$ | 2 | Rev-Erb-inhibition |
| $p$ | 6 | Ror-activation |
| $q$ | 3 | Ror-inhibition |
| $n$ | 2 | Bmal-activation |
| $m$ | 5 | Bmal-inhibition |
| $y_{10}$ | 0*omega | Per |
| $y_{20}$ | 0*omega | Cry |
| $y_{30}$ | 0*omega | Rev-Erb |
| $y_{40}$ | 0*omega | Ror |
| $y_{50}$ | 0*omega | Bmal |
| $d_{z1}$ | 0.23 | CRYC |
| $d_{z2}$ | 0.25 | PERC |
| $d_{z3}$ | 0.6 | PERC* |
| $d_{z4}$ | 0.2 | PERC*/CRYC |
| $d_{z5}$ | 0.2 | PERC/CRYC |
| $d_{z6}$ | 0.31 | REV-ERBC |
| $d_{z7}$ | 0.3 | RORC |
| $d_{z8}$ | 0.73 | BMALC |
| $d_{y1}$ | 0.3 | Per |
| $d_{y2}$ | 0.2 | Cry |
| $d_{y3}$ | 2 | Rev-Erb |
| $d_{y4}$ | 0.2 | Ror |
| $d_{y5}$ | 1.6 | Bmal |
| $d_{x1}$ | 0.08 | CLOCK/BMAL ** -23 PD Huge Phase deriv -38 PD |
| $d_{x2}$ | 0.06 | PER*N/CRYN ** |
| $d_{x3}$ | | PERN/CRYN |
| $d_{x5}$ | 0.17 | REV-ERBN |
| $d_{x6}$ | 0.12 | RORN ** -28 PD |
| $d_{x7}$ | 0.15 | BMALN |
| $k_{f_{x1}}$ | 2.3 | CLOCK/BMAL-complex formation [hour-1] |
| $k_{d_{x1}}$ | 0.01 | CLOCK/BMAL-complex dissociation [hour-1] |
| $k_{f_{z4}}$ | 1/omega | PERC*/CRYC-complex formation [(a.u.×hour)-1] |
| $k_{d_{z4}}$ | 1 | PERC*/CRYC-complex dissociation [hour-1] |
| $k_{f_{z5}}$ | 1/omega | PERC/CRYC-complex formation [(a.u.×hour)-1] |
| $k_{d_{z54}}$ | 1 | PERC/CRYC-complex dissociation [hour-1] |
| $k_{i_{z4}}$ | 0.2 | PERC*/CRYC |
| $k_{i_{z5}}$ | 0.1 | PERC/CRYC |
| $k_{i_{z6}}$ | 0.5 | REV-ERBC |
| $k_{i_{z7}}$ | 0.1 | RORC ** 12.9 PD |
| $k_{i_{z8}}$ | 0.1 | BMALC ** 15.6 PD |
| $k_{e_{x2}}$ | 0.02 | PER*N/CRYN ** -21 PD |
| $k_{e_{x3}}$ | 0.02 | PERN/CRYN |
| $k_{ph_{z2}}$ | 2 | PERC-phosphorylation rate |
| $k_{d_{ph_{z3}}}$ | 0.05 | PERC*-dephosphorylation rate |
| $k_{p1}$ | 0.4 | PERC |
| $k_{p2}$ | 0.26 | CRYC |
| $k_{p3}$ | 0.37 | REV-ERBC |
| $k_{p4}$ | 0.76 | RORC |
| $k_{p5}$ | 1.21 | BMALC |
| $k_{t1}$ | 3*omega | Per-activation rate |
| $k_{i1}$ | 0.9*omega | Per-inhibition rate |
| $k_{t2}$ | 2.4*omega | Cry-activation rate PLOSGEN PERTUR 5.3 TO SIM RAS |
| $k_{i2}$ | 0.7*omega | Cry-inhibition rate |
| $k_{i21}$ | 5.2*omega | Cry-inhibition rate |
| $k_{t3}$ | 2.07*omega | Rev-Erb-activation rate |
| $k_{i3}$ | 3.3*omega | Rev-Erb-inhibition rate |
| $k_{t4}$ | 0.9*omega | Ror-activation rate |
| $k_{i4}$ | 0.4*omega | Ror-inhibition rate **15.5 PD |
| $k_{t5}$ | 8.35*omega | Bmal-activation rate |
| $k_{i5}$ | 1.94*omega | Bmal-inhibition rate |
| $V_{1max}$ | 1*omega | Per |
| $V_{2max}$ | 2.92*omega | Cry |
| $V_{3max}$ | 1.9*omega | Rev-Erb |
| $V_{4max}$ | 10.9*omega | Ror |
| $V_{5max}$ | 1*omega | Bmal |

### S13 Previous time-telling algorithms

A summary of the existing time-telling or clock dysfunction measuring studies is given in this section. All methods are for calculating time (internal or external) in some way from transcriptomes, except for ΔCCD which attempts to calculate clock dysfunction from whole datasets (not single samples as TimeTeller’s Θ can).

The key difference between TimeTeller and the other methods is TimeTeller’s use of the covariance structure of an estimate of *P* (*g*|*t*) where *g* = (*g*_1_, …, *g*_*G*_) is a vector of the expressions of *G* genes in WT samples and *t* is time. We explain in Note S4 why the distribution *P* (*t*|*g*) (related to *P* (*g*|*t*) by Bayes theorem) is the appropriate likelihood to study to determine clock precision and why the variance and confidence limits of the maximum likelihood estimate using *P*(*t*|*g*) depends crucially on this covariance structure.

#### S13.1 Molecular timetabling method (MTTM)

The aim of the study in Ueda et al. (*25*) was to detect individual body time (BT) via a single-time-point assay so that BT information could be exploited to optimize medication strategies. Although this study uses BT and not clock time in peripheral tissue in their estimation, the results in the study show that in controlled mouse experiments, BT and CT are equivalent. To detect time-indicating genes, i.e. genes whose expression exhibits circadian rhythmicity with high amplitude, the expression profile of each gene was first analysed for rhythmicity and amplitude. The top *N* rhythmic genes were then chosen for use in the timetabling method. They normalised the expression profile of each gene using its mean and standard deviation in the molecular timetable. To make the estimation for the time of a single sample, a single set of gene expressions of unknown time, is normalised with the same means and standard deviations as the training set. Cosines of 10 minute resolution are fitted to these single time points and the best BT estimate is given by the time with the best correlation.

#### S13.2 ZeitZeiger

ZeitZeiger (Hughey et al. (*26*)) ia a supervised learning method for high dimensional data from an oscillatory system. ZeitZeiger is conceptually similar to supervised principal components (SPC (*27*)) as the genes used in the principal component analysis are chosen so that they show maximum variation over time. There are 2 main parameters that need to be chosen; sumabsv controls how many genes form each sparse PC, and nSPC determines how many SPCs are used for prediction.

After suitable data preparation (e.g. batch correction) for each feature (e.g. gene), *j* = 1, …, *p*, ZeitZeiger estimates the time-dependent mean, *f*_*j*_(*t*), by fitting a periodic smoothing spline to the training observations *g*_*ij*_ of the expression of gene *j* in sample *i*. It estimates the variance 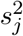 of the expression of gene *j* about this mean curve. The data matrix *X* has as its *i*th row the expression vector (*g*_*ij*_). These splines are discretised in time (*m* equally spaced times) mean-zeroed and scaled using the standard deviation *s*_*j*_. The resulting vectors form the columns of a *m* × *p* matrix *Z*. The right singular vectors from the penalized matrix decomposition of *Z* are the sparse principal components (SPCs) that are used in the analysis.

The SPCs are used to project the training data from high-dimensional feature-space to low-dimensional (typically 2d) SPC-space. Each data vector *g* = (*g*_1_, …, *g*_*p*_) is projected to *ω*_*k*_ = *ω*_*k*_(*g*) = *g* · *V*_*k*_. For the *k*th SPC, *ω*_*k*_, the time dependent mean 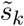, and the variance 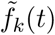 about this mean are calculated as *g* varies over the training data in *X*.

To analyse an independent sample *g* one projects *g* to obtain *ω* = *gV* and then forms the lkelihoods

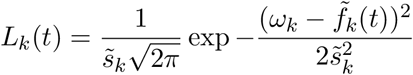

combining them as 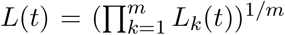. The time *T* is estimated as the time at which *L*(*t*) is maximal.

Because of batch and similar effects one has to combine training and test data and use batch removal techniques and software such as ComBat to perform cross-study and cross-organ normalization before applying ZeitZeiger.

We compare ZeitZeiger and TimeTeller likelihood curves and the associated Θs in Note S10.4.

#### S13.3 BodyTime as a diagnostic clinical tool

Wittenbrink *et al.* (*28*) developed a strategy that estimates internal body time from a single blood sample, in order to inform personalised medicine strategies.

Firstly, they created a novel human timecourse data set using RNAseq of blood monocytes of 12 males (14 samples every 3 hours over 40 hours). They searched for circadian biomarkers i.e. for genes that have “time telling properties” and subsequently acquired nanostring data for these 34 genes.

After doing RNAseq-nanostring platform comparison, they concluded that there was a more than satisfactory performance of the candidate genes as time telling genes using the nanostring platform. They applied ZeitZeiger to this nanostring data set and validated with a leave-one-subject out approach. An external data set (VALI) was used as an external validation, which resulted in accurate predictions of body time.

#### S13.4 BIO CLOCK

The paper (*29*) has two themes; one of period detection and the other of estimating time from single samples. The methods for BIO CLOCK are simply described as a supervised deep learning algorithm using neural networks, with few method specifics discussed.

#### S13.5 CYCLOPS

Anafi et al. (*30*) published CYCLOPS : a neural network that finds circadian patterns in datasets (*31*). This study does not present an algorithm that can (yet) tell the time of single samples (and does not claim to be able to), but it has clear value in this field. The study uses a (quasi) unsupervised machine learning algorithm (an autoencoder) to construct a cyclic periodic timecourse, using unordered large datasets of expression measurements according to the clock genes, by mapping their eigengenes to ellipses. These known rhythmic genes are given as prior information. The number of eigengenes retained was set so that > 85% of variance was captured. CYCLOPS optimally weights and combines the eigengenes, in order to create the closest thing to an ellipse. These points are subsequently ordered along that ellipse using a neural network. Optimal weighting and combination was performed through use of a circular node autoencoder.

#### S13.6 PLSR

Laing et al. (*32*) published a study that uses a partial least squares regression method to estimate melatonin cycles from transcriptome data. They used a large, novel human transcriptome dataset, which consisted of timecourses of melatonin and gene expression in sleep deprived states. The study is not strictly speaking presenting a “timetelling” model. The authors instead attempt to predict melatonin phase from the transcriptome of blood samples from human volunteers. There is no new method developed in this study, instead a new application of an existing and established supervised ML method, Partial Least Squares Regression (*33*).

#### S13.7 ΔCCD

Shilts et al. (*34*) developed the ΔCCD clock coefficient of dysfunction. Using 12 clock genes, they examine the (Spearman) correlations between all pairs of these genes and use these correlations as the “healthy” standard. Spearman’s rank is used as a measure of correlation of two genes, denoted by the measure *ρ*. These *ρ* coefficient tables are produced for independent data sets. The ΔCCD metric is a simple euclidean distance metric that is a measure of distance between the 12 × 12 standard table, and the 12 × 12 test table. This metric is similar to the aims of the Θ metric from our TimeTeller, but ΔCCD is calculated over a whole set of samples (multiple points are needed to calculate a spearman’s correlation), whereas Θ can be calculated for single time point samples.

## Footnotes

1 The Research Ethics Board at Sunnybrook Health Science Centre approved the clinical protocol for this study.

2 Unfortunately, a different (custom) microarray was used to assess the tumour samples, which excluded the majority of the clock genes.

3 Throughout these notes. denotes the transpose of a matrix.

